# Mesoscopic in vivo human T_2_* dataset acquired using quantitative MRI at 7 Tesla

**DOI:** 10.1101/2021.11.25.470023

**Authors:** Omer Faruk Gulban, Saskia Bollmann, Renzo Huber, Konrad Wagstyl, Rainer Goebel, Benedikt A. Poser, Kendrick Kay, Dimo Ivanov

**Author notes:** Equal contribution.

## Abstract

Mesoscopic (0.1-0.5 mm) interrogation of the living human brain is critical for advancing neuroscience and bridging the resolution gap with animal models. Despite the variety of MRI contrasts measured in recent years at the mesoscopic scale, in vivo quantitative imaging of T_2_* has not been performed. Here we provide a dataset containing empirical T_2_* measurements acquired at 0.35 × 0.35 × 0.35 mm^3^ voxel resolution using 7 Tesla MRI. To demonstrate unique features and high quality of this dataset, we generate flat map visualizations that reveal fine-scale cortical substructures such as layers and vessels, and we report quantitative depth-dependent T_2_* (as well as R_2_*) values in primary visual cortex and auditory cortex that are highly consistent across subjects. This dataset is freely available at https://doi.org/10.17605/OSF.IO/N5BJ7, and may prove useful for anatomical investigations of the human brain, as well as for improving our understanding of the basis of the T_2_* -weighted (f)MRI signal.

## 1 Introduction

A central goal in neuroscience is to achieve a comprehensive assessment of the structure of the living human brain. Centuries of work have been dedicated to characterizing the human brain from its macroscopic features (e.g., lobes, gyri, sulci) to its microscopic constituents (e.g., dendrites, cell bodies) (Finger et al., 2009, Ch. 1-10). The history of neuroscience has witnessed steady improvements in technologies for imaging the brain at increasing levels of detail (Bentivoglio & Mazzarello, 2009). From the late 19^th^ to the early 20^th^ century, major advancements in tissue staining and photography opened new doors for measuring the human brain’s cytoarchitecture (Geyer, 2013, Part 1), myeloarchitecture (Nieuwenhuys, 2013), fiber architecture (Kleinnijenhuis, 2014, Ch. 1), and angioarchitecture (Pfeifer, 1940).

In seminal works of neuroscience, it has been acknowledged that post mortem experiments on animal and human brains are valuable but temporary necessities (Stahnisch, 2010). Fortunately, new technologies that enable study of the living human brain have emerged. The second half of the 20^th^ century has brought computerized tomography (CT), positron emission tomography (PET), and magnetic resonance imaging (MRI) (Raichle, 2009a). Though these new technologies were far from the microscopic details provided by older methods, they quickly became indispensable due to their non-invasive nature. MRI has proven particularly useful for its flexibility to measure different tissue contrasts as well as hemodynamic measures of brain activity (Raichle, 2009b). Recently, MRI at ultra-high magnetic field strength (≥7 Tesla) has breached the mesoscopic scale, which we define as resolutions between 0.1 mm and 0.5 mm (Edwards et al., 2018). Particularly, openly accessible datasets by Bollmann et al. (2022), Federau and Gallichan (2016), Lüsebrink et al. (2021), and Lüsebrink et al. (2017), Schira et al. (2022) have facilitated the incorporation of mesoscopic MRI into the toolkit of practicing neuroscientists. However, despite the rich variety of MRI contrasts acquired by the previous work (Barbier et al., 2002; Bollmann et al., 2022; Bridge et al., 2005; Budde et al., 2011; Dinse et al., 2015; Duyn et al., 2007; Federau & Gallichan, 2016; Fracasso et al., 2016; Kemper et al., 2018; Lüsebrink et al., 2021; Lüsebrink et al., 2017; Mattern et al., 2018; Petridou et al., 2010; Sanchez Panchuelo et al., 2021a; Sánchez-Panchuelo et al., 2012; Schira et al., 2022; Tardif et al., 2015; Trampel et al., 2011; Turner et al., 2008; Zwanenburg et al., 2011), there has been no openly accessible quantitative in vivo human T_2_* measurements at mesoscopic resolution (see **Supplementary Table 1)**.

In this paper, we deliver this missing element: quantitative in vivo human T_2_* measurements. The data are acquired at the challenging resolution of 0.35 × 0.35 × 0.35 mm^3^ voxels at 7 Tesla and cover a substantial portion (approximately one-third) of the human brain, primarily including the visual and auditory cortices. To highlight the unique features and high quality of this dataset, (i) we provide a local (voxel-wise) qualitative assessment of cortical substructures using three dimensional flat maps, highlighting cortical substructures such as layers and vessels; and (ii) we provide a global (region-of-interest-wise) quantitative report of depth-dependent (also referred as layer-specific or laminar) T_2_* values of the cortical gray matter, superficial white matter, and above pial surface at the primary visual and auditory cortices. The main contribution of this paper is to demonstrate a practicable data acquisition protocol for collecting mesoscopic anatomical images in living humans, and to deliver a novel, freely available dataset for further use by the community. We believe this dataset will be highly useful for development of novel biomarkers of cortical laminar structure (e.g. McColgan et al., 2021), for improving the parcellation of cortical areas (e.g. Gulban et al., 2020), and for better understanding the basis of layer-fMRI signals (e.g. Havlicek and Uludag, 2020).

## 2 Methods

### 2.1 Participants

Five healthy participants (2 females, 27-35 years old) were recruited for the study. These participants were selected for their experience in participating in 7 T experiments and for their ability to keep their heads still for a long time. Similar preselection strategies were previously used in high resolution MRI studies (Allen et al., 2022; Bollmann et al., 2022; Tardif et al., 2015). Informed consent was obtained from each participant before conducting the experiment. The study was approved by the research ethics committee of the Faculty of Psychology and Neuroscience of Maastricht University and experimental procedures followed the principles expressed in the Declaration of Helsinki.

### 2.2 Data Acquisition

#### 2.2.1 Session 1: T_2_* with ME GRE

In the first session, we acquired 3D multi-echo gradient recalled echo (ME GRE) images with bipolar readouts to measure T_2_* (see **Table 1)** using 7T MRI (Siemens) with a 32-channel Rx head coil (Nova). More specifically, our ME GRE sequence was a customized version of the vendor’s product sequence (multi-echo FLASH sequence with ASPIRE coil combination developed by Eckstein et al., 2018). The nominal image resolution was 0.35 × 0.35 × 0.35 mm^3^, and parameters included TR = 30 ms, TE_1-6_ = [3.83, 8.20, 12.57, 16.94, 21.31, 25.68] ms, FA = 11°, 576 × 576 × 104 voxels, FOV = 20.16 × 20.16 × 3.64 cm^3^, GRAPPA = 2, elliptic k-space filling, 15 minutes duration, and no flow compensation. See **Figure 1** for the positioning of our imaging slab. Four successful ME GRE runs totalling 60 minutes were acquired in this scanning session. Between acquisitions of the 3D ME GRE we changed the phase-encoding direction by 90° (right-left, anterior-posterior, left-right, posterior-anterior). These 90° changes were introduced in order to control the direction of the blood motion artifact which is caused by the time difference between the phase encoding and readout stages (this artifact is called “spatial misregistration of the vascular flow” by Larson et al., 1990). As a result of this blood motion artifact, we see the vector component of the blood flow in the readout direction as a displacement (see **Supplementary Figure 1)**. Additionally, the bipolar readouts have odd- and even-numbered echos acquired with opposite (180°) readout directions, which is known to introduce a slight spatial shift between their corresponding images due to the change in readout direction (Cohen-Adad, 2014). However, we found that this shift is negligible compared to the blood motion artifact. We have also acquired a whole-brain magnetization prepared 2 rapid acquisition gradient echos (MP2RAGE) image (Marques et al., 2010) at 0.7 mm isotropic resolution. Parameters of this whole-brain MP2RAGE acquisition was TR = 5000 ms, TI_1-2_ = [900, 2750] ms, TE = 2.46 ms, FA_1-2_ = [5°, 3°], 320 × 320 × 240 voxels, FOV = 22.4 × 22.4 × 16.8 cm^3^. The shimming procedure included the vendor-provided routines to maximize the field homogeneity within the imaging slab. We found that the shimming was quite effective (spatially varying off-resonance frequencies occur on a very large spatial scale).

**Table 1:** Acquisition parameters for the high-resolution scanning protocols (7 T MRI).

|  |  |  |
| --- | --- | --- |
| Sequence | ME GRE | MP2RAGE |
| Voxel size | $0.35 \times 0.35 \times 0.35 \text{ mm}^3$ | $0.35 \times 0.35 \times 0.35 \text{ mm}^3$ |
| Field of view (FOV) | $201.6 \times 201.6 \times 36.4 \text{ mm}^3$ | $201.6 \times 201.6 \times 42 \text{ mm}^3$ |
| Slab dimensions | $576 \times 576 \times 104 \text{ voxels}$ | $576 \times 576 \times 120 \text{ voxels}$ |
| Bandwidth | 240 Hz/Px | 250 Hz/Px |
| TR | 30 ms | 5000 ms |
| TE | [3.83, 8.20, 12.57, 16.94, 21.31, 25.68] ms | 2.91 ms |
| Flip angle (FA) | $11^\circ$ | $[4^\circ, 5^\circ]$ |
| TI | N/A | [800, 2700] ms |
| Read-out gradient mode | Bipolar | Monopolar |
| Phase partial Fourier | Off | 6/8 |
| GRAPPA | 2 | 2 |
| Volume acquisition time | 15 min | 10 min |
| Total duration | $4 \times 15 \text{ min} = 60 \text{ min}$ | $6 \times 10 \text{ min} = 60 \text{ min}$ |

**Figure 1:**
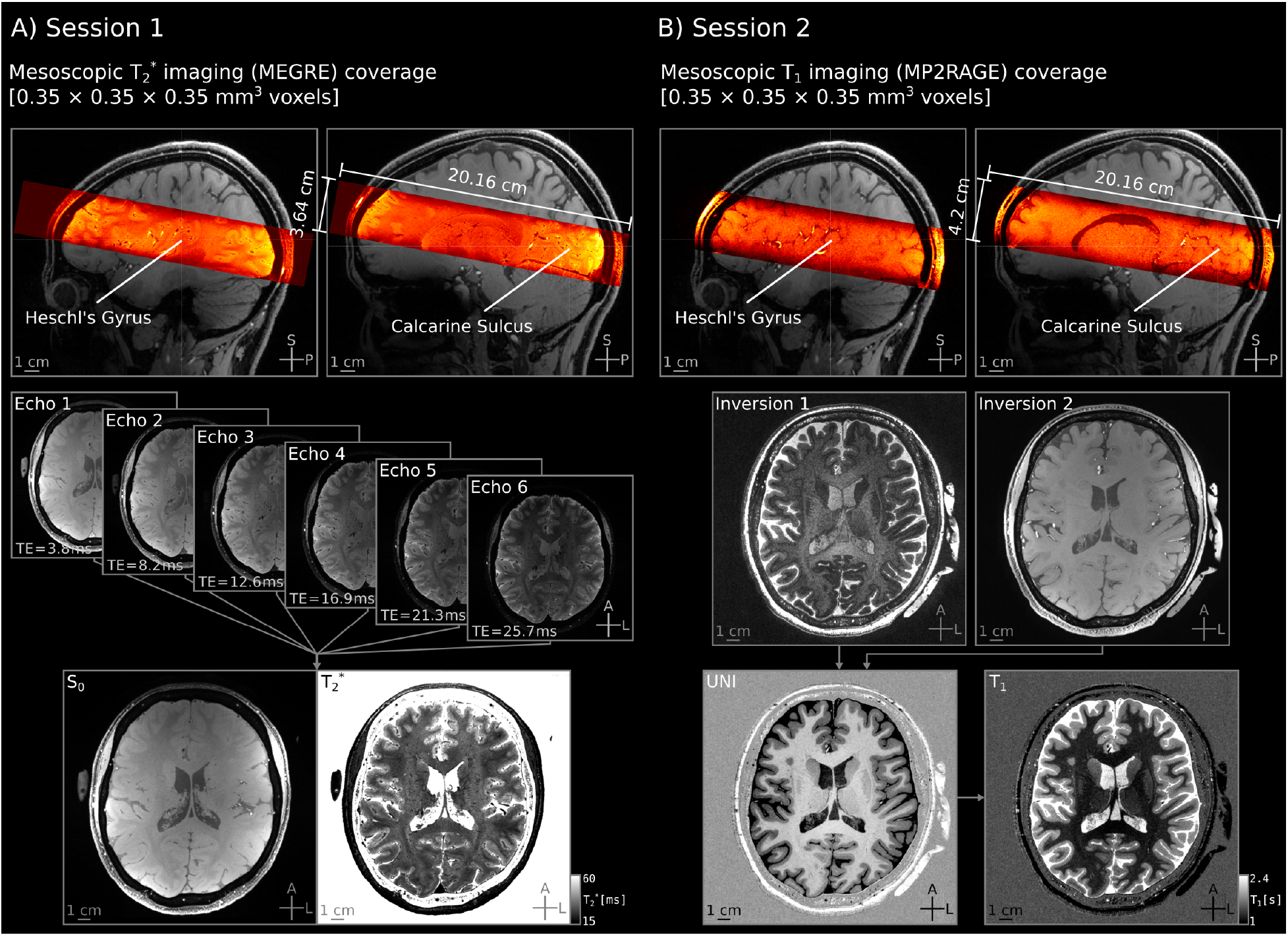
Spatial coverage and image quality of the MRI data acquisition. Here we show data quality for an example participant (sub-01) for the first **(A)** and second scan sessions **(B)** after preprocessing. High-resolution imaging slabs (in warm colors) are overlaid on top of the whole-brain lower-resolution image. The imaging slabs are positioned to cover both the calcarine sulci and Heschl’s gyri (transverse temporal gyrus). Transversal images are averaged across runs. Despite the very high spatial resolution (0.35 mm isotropic), signal-to-noise ratio of our averaged images is high, clearly showing fine details in each individual (see **Supplementary Figure 2)** and the signal-to-noise ratio of individual runs are consistent across acquisitions (see **Supplementary Figures 3–4)**. See **Table 1** for acquisition parameters, and **Section 2.7** for data availability and sequence parameter documents.

#### 2.2.2 Session 2: T_1_ with MP2RAGE

In the second session, we acquired mesoscopic MP2RAGE images (Marques et al., 2010) (see **Table 1)** using the same setup. These T_1_ data were acquired primarily for cortical gray matter segmentation purposes and were not intended to serve as quantitative T_1_ measurement. The nominal image resolution was equal to the T_2_* images at 0.35 × 0.35 × 0.35 mm^3^, and parameters included TR = 5000 ms, TE = 2.91 ms, TI_1-2_ = [800, 2700] ms, FA_1-2_ = [4°, 5°], 576 × 576 × 120 voxels, 20.16 × 20.16 × 4.2 cm^3^ slab dimensions, GRAPPA = 2, Partial Fourier = 6/8, Bandwidth = 250 Hz/Px, and 10 minutes duration. See **Figure 1** for the positioning of our imaging slab. Six successful MP2RAGE runs totalling 60 minutes were acquired in this scanning session. For completeness, we changed the phase-encoding direction by 90° in each run similar to the ME GRE images acquired in Session 1. However, the effect of the blood motion artifact was negligible because of the short TE used in MP2RAGE images compared to ME GRE. Our MP2RAGE sequence was a customized version of the vendor’s product sequence. We hard-coded the bandwidth time product of the excitation pulse parameter to 25, which increased the sharpness of the slab.The shimming procedure included the vendor-provided routines to maximize the field homogeneity within the imaging slab. We found that the shimming was quite effective as in our first session.

### 2.3 Data Analysis

Data processing and analysis scripts referenced in this section are available at: https://github.com/ofgulban/meso-MRI (v1.0.2 saved at https://zenodo.org/record/7210802). See **Supplementary Figures 14 and 15** for an overview of our processing pipeline.

#### 2.3.1 T_2_* images (ME GRE)

Each ME GRE image series consists of odd- and even-numbered echos with opposite readout direction (while phase-encoding was rotated by 90° between acquisitions). Our processing pipeline aimed at reducing two sources of artifacts while improving the signal-to-noise ratio: (i) head motion across acquisitions, (ii) blood motion artifacts across echos. The processed data is then used to compute T_2_* and S_0_ values. Note that we did not need to deface our images as face region is not included in our imaging slab to begin with. See **Supplementary Figure 2** for an overview of the preprocessed data quality across participants which shows that the fine detailed cortical substructures are captured consistently across all participants.

#### 2.3.2 T_2_* head motion correction

There are two types of bulk head motion to consider: (i) “within-run head motion” which introduces blur and ringing artifacts into the images, and (ii) “in between runs” head motion which blurs the average of images. To address the within-run head motion, we preselected participants who are experienced with being scanned and with minimizing head motion. This strategy has been used successfully in Allen et al. (2022), Bollmann et al. (2022), and Tardif et al. (2015). In addition, we kept our run durations between 10-15 minutes. Durations above 20 minutes have been shown to require prospective motion correction (Lüsebrink et al., 2017; Mattern et al., 2018; Stucht et al., 2015), or the use of fat-based motion navigators (Federau & Gallichan, 2016; Gallichan et al., 2016). However, shorter acquisition durations together with participant preselection has been shown to generate similar quality data up to 0.16 mm isotropic resolution (Bollmann et al., 2022). While Bollmann et al. (2022) reports approximately 1 out of 4 runs had to be discarded, we did not discard any data upon quality controlling our images. An overview of our unpreprocessed data in **Supplementary Figure 3** shows the effectiveness of our participant preselection strategy yielding high quality images that are not strongly affected by within run head motion in any of the individual runs.

In what follows, we describe the processing steps to address the “in between runs” head motion. We started by cropping our images to exclude the frontal brain regions (`01_crop.py`). We note that this cropping is done to reduce computational requirements (RAM usage) for the upcoming steps. Then we averaged signal intensities across all echos per voxel (`02_avg_echos.py`). This averaged image improves signal-to-noise ratio and is used only in the estimation of motion. We also upsampled both the original echos and the averaged images to 0.175 mm isotropic resolution with cubic interpolation; this allows fine-scale detail to be preserved during the motion correction process (Allen et al., 2022) (`03_upsample.py`). Head motion was estimated from the averaged images using rigid-body transformation (6 degrees of freedom) while using a manually defined brain mask covering our regions of interest; then, data were corrected using linear interpolation (`04_motion_correct.py`) (Yushkevich et al., 2006). Note that all runs are co-registered to the first ME GRE run. The estimated rigid body transformation matrices were applied to each upsampled echo separately (`05_split_echos.py, 07_apply_reg.py, 08_merge_echos.py`), resulting in a final set of brain volumes with nominal 0.175 mm isotropic resolution.

#### 2.3.3 T_2_* blood motion artifact mitigation

After motion correction, we averaged the ME GRE images with the same phase-encoding axis for each echo (PE_x_ for right-left and left-right; PE_y_ for anterior-posterior and posterior-anterior) to improve signal-to-noise ratio (`09_average_same_PE_axes.py`). This step is performed because the blood motion artifact direction is independent of the *phase-encoding direction*, and only dependent on the *vector component of the blood flow along the readout axis* (Larson et al., 1990). Then we composited a new image by selecting voxels from the PE_x_ and PE_y_ phase encoding axis images using a minimum operator (`10_composite.py`). Selecting the minimum intensity observed across the two images mitigates the blood motion artifact because the blood motion artifact occurs as a positive term added to the local signal. Since the blood motion artifact will be different across images with 90° rotated phase-encoding axes (PE_x_ and PE_y_), it is conceivable to composite a new image (i.e. a new set of echos) by selecting the signal of each voxel from the original set that is not affected by the artifact (a voxel affected by the artifact in PE_x_ highly unlikely to be affected in PE_y_). Such image compositing operations are commonly used in the movie-making visual effects field (Brinkmann, 2008). In compositing, the main idea is to piece together a new image by using parts of multiple other images (see **Supplementary Figure 1)**.

As the last step to mitigate the residual blood motion artifacts, we detected voxels that do not decay across echos (e.g. points on the decay curve showing a higher signal compared to the echo that comes before), and replaced its value with an average of the echo before and after (`11_fix_nondecay.py`). This fix for non-decaying voxels is performed in order to eliminate residual blood motion outliers that could affect the T_2_* fit. In our experience, this fix is needed only for a tiny portion of voxels. This situation is very unlikely to occur after our previous compositing step; however, when it does occur, it is due to the highly curved and convoluted vessel configuration in the region. Future work could explore rotating the phase encoding axis in 45° increments or around another axis to increase the chances of acquiring data that are not affected by the blood motion artifact.

Note that our procedure is best viewed as *mitigating* as opposed to *correcting* the blood motion artifact: we are attempting to suppress the misencoded artery signal, not move it back to its correct location. We also highlight that we used the minimum operator as the simplest way of mitigating the blood motion artifact. Alternative artifact-mitigation strategies that might more efficiently combine the data constitute an interesting direction for future work.

#### 2.3.4 T_2_* fitting

We fit a monoexponential decay function 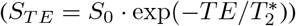 to the processed ME GRE images by fitting a line to the logarithm of the signal using ordinary least-squares (`12_fit_T2star.py`) (Cohen-Adad et al., 2012) using Nibabel, Scipy, and Numpy (Brett et al., 2017; Jones et al., 2001; Van Der Walt et al., 2011). Note that in addition to providing an estimate of T_2_*, this procedure also provides an estimate of S_0_ (see **Figure 1)**.

#### 2.3.5 T_1_ images (MP2RAGE)

Our processing strategy for the T_1_ images is aimed at correcting head motion across acquisitions while boosting signal-to-noise ratio through averaging. Note that we did not need to deface our images as face region is not included in our coverage to begin with. See **Supplementary Figure 4** for an overview of the preprocessed data quality across participants which shows that we have attained a good contrast to delineate the inner and outer boundaries of the cortical gray matter.

#### 2.3.6 T_1_ head motion correction

No MP2RAGE data was discarded due to within-run head motion. An overview of our unprocessed data in **Supplementary Figure 4** shows that our participant preselection strategy was effective to mitigate the within-run head motion artifacts, similar to the T_2_* (ME GRE) imaging shown in **Supplementary Figure 3**. We concluded that our MP2RAGE images were of good quality and were ready for coregistration across runs followed by averaging to boost the signal-to-noise ratio.

MP2RAGE images were processed similarly to the ME GRE images for the first two steps of cropping and upsampling (`01_crop.py, 02_upsample.py`). All runs are co-registered to the first MP2RAGE run using the second inversion time (INV2) contrast with the GREEDY (Yushkevich et al., 2016) registration algorithm (acquired from: https://github.com/pyushkevich/greedy). We used the second inversion time contrast to drive registration because it has the best overall brain and non-brain signal-to-noise ratio. A brain mask focusing on our regions of interest was manually drawn in ITK-SNAP (Yushkevich et al., 2006) and used to constrain the registration cost metric. Registration was estimated using GREEDY rigid-body transformation (6 degrees of freedom) and data were corrected using linear interpolation (`03_motion_correct.py`). The estimated transformations were applied to all of the MP2RAGE contrasts (`04_apply_reg.py`). Finally, we computed the average across all motion-corrected images (`05_average.py`).

#### 2.3.7 T_1_ registration to T_2_* images

The MP2RAGE data were registered to the ME GRE using the second inversion time (INV2) contrast from the MP2RAGE and the S_0_ image resulting from T_2_* fitting of the ME GRE. This choice was made due to the similarity of tissue contrast in these images. We used the GREEDY registration with 6 degrees of freedom and linear interpolation (`06_register_to_T2s.py`).

### 2.4 Segmentation

Accurate identification of gray and white matter is critical for proper interpretation of cortical architecture. We carefully segmented four main regions of interest in the registered MP2RAGE UNI images. These regions consisted of brain tissue in and around the calcarine sulcus and Heschl’s gyrus in each hemisphere. To determine these regions, we centered a spherical mask at each of the relevant sulcal and gyral landmarks. We refer to these masks as “scoops of interest” (see **Figure 2)**. We used an interactive intensity histogram thresholding method (Gulban et al., 2018) to obtain our initial tissue segmentation within each scoop of interest based on the MP2RAGE UNI contrast. After this step, each scoop of interest was manually edited by an expert (O.F.G.) and quality controlled for accurate and precise tissue segmentation (R.H.). The manual editing process took approximately 8 to 10 hours for each subject. We have chosen this laborious manual segmentation process over a fully automatic one because of the lack of optimized and validated segmentation tools for our very high resolution data. The resulting tissue segmentations are available as a part of our data repository and can be freely inspected or used for validating the results of automatic algorithms (Bazin et al., 2014; Svanera et al., 2021). Note that we used the scoops of interest rather than segmenting all of the brain tissue available within our imaging slabs to focus our efforts on achieving the best segmentation for our regions of interest.

**Figure 2:**
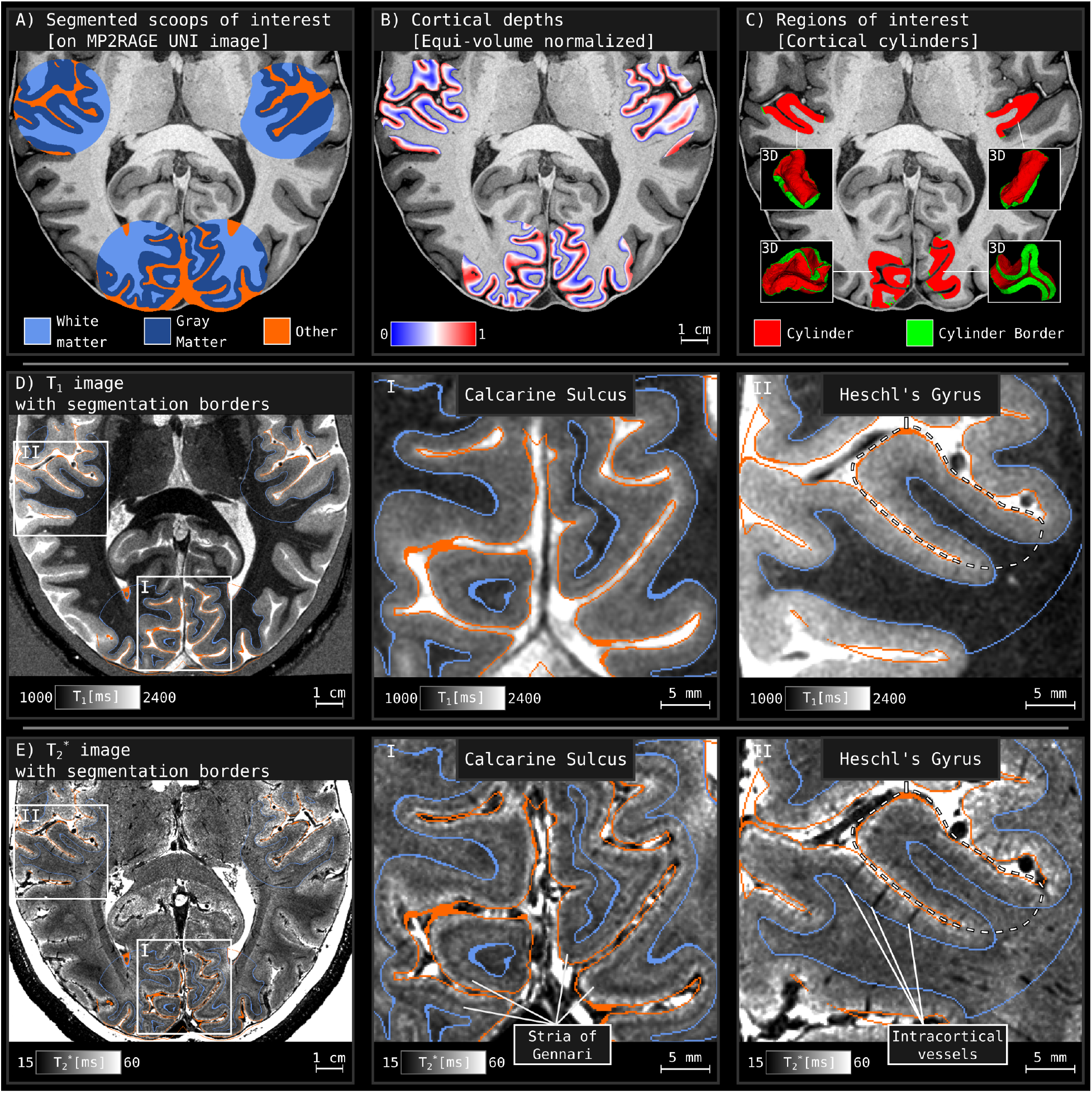
Segmentation, cortical depths, and regions of interest identified for an example subject (sub–04). **(A-C)** Major processing steps for characterizing our cortical regions of interest. The cortical cylinders in C were geodesically centered at the calcarine sulci and Heschl’s gyri anatomical landmarks based on the middle gray-matter voxels. Careful manual tissue segmentation was performed, enabling extraction of gray-matter boundaries and cortical surface topology. **(D–E)** T_1_ and T_2_* measurements in the regions of interest with overlaid gray-matter boundaries. Notice that the quality of the T_2_* image allows visualization of fine-scale cortical substructures such as the Stria of Gennari and intracortical vessels in the zoomed-in panels **I-II**. Detailed images of the other participants can be seen in **Supplementary Figure 2**.

### 2.5 Cortical Depths

After completing the tissue segmentation, we used LayNii v.2.2.0 (Huber et al., 2021) to compute equi-volume cortical depths (Bok, 1959). Specifically, we prepared the segmentation input, i.e. the motion-corrected and averaged T_1_ -weighted images (`00_prep.py`), and used the LN2_LAYERS program to compute equi-volume normalized cortical depth measurements for each gray matter voxel (`01_layers.py`). Note that the voxel-wise cortical depth metric computed in LN2_LAYERS ranges between 0 and 1 and reflect normalized units: cortical depth measurements (in mm) are normalized by local cortical thickness measurements (in mm) and then adjusted within this closed vector space to determine the equi-volume metric (see **Figure 2B** and **Figure 3B**) (this implementation is described in Huber et al., 2021). In addition, this program computes cortical curvature at each gray-matter voxel. See (Bok, 1959; Waehnert et al., 2014) for general references on the equi-volume principle of cortical layering.

**Figure 3:**
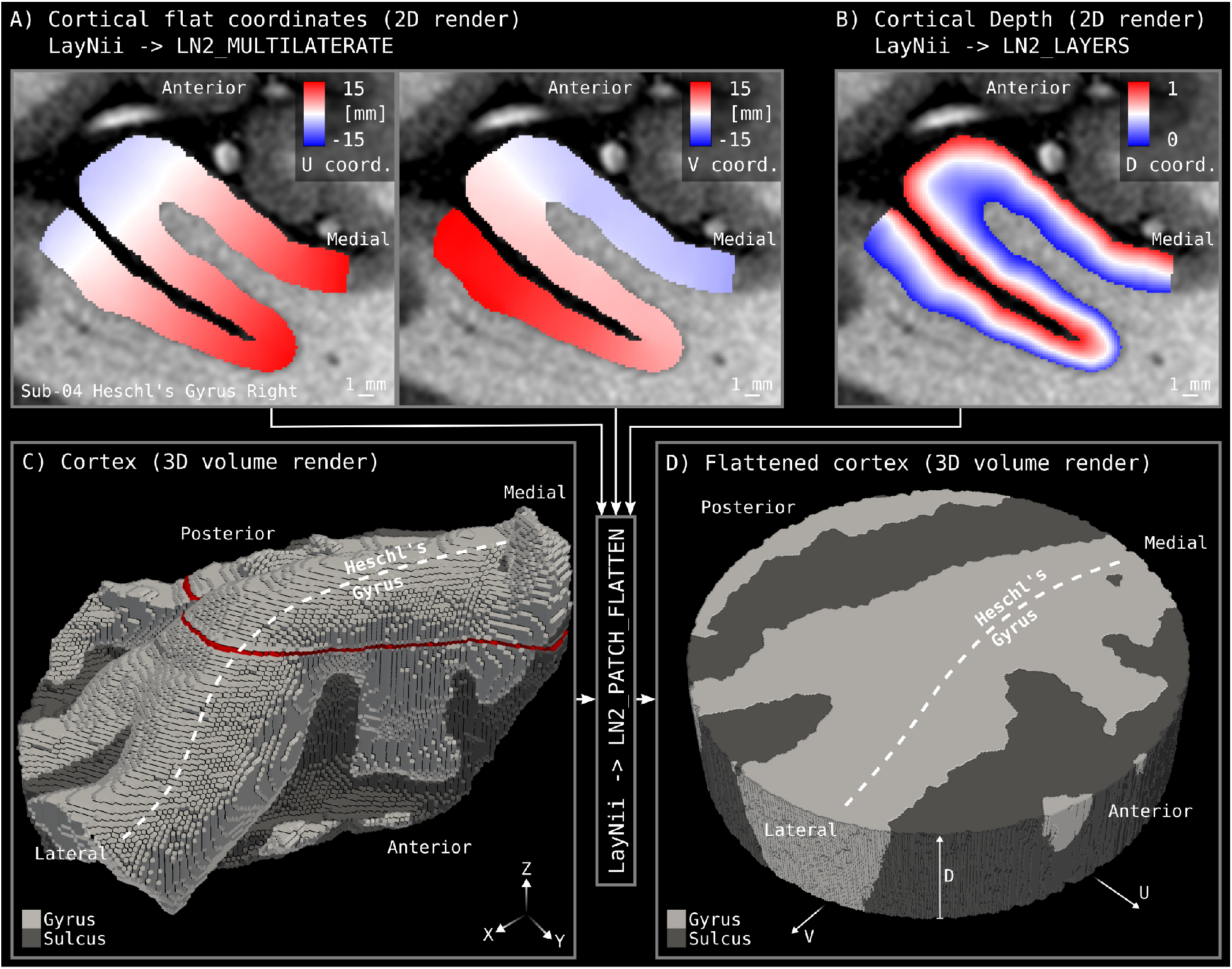
3D to 3D cortical flattening. Cortical cylinders in the form of *virtual Petri dishes* are used for flattening our regions of interest. A cylinder is centered at our anatomical landmarks and its radius is set to 15 mm. The cylinder height covers the whole cortical thickness. **(A)** Cortical flat coordinates visualized for a 2D slice. **(B)** Cortical equi-volume depth assignment for the same slice. **(C)** Structure of cortical cylinder in original 3D folded brain space (XYZ coordinates). The red line indicates the location of the 2D slice in panels A and B. **(D)** Structure of cortical cylinder after flattening (UVD coordinates).

To facilitate anatomical quantification, we also calculated distances for voxels that lie beyond the inner and outer gray-matter boundaries. This was done using LayNii program LN2_GEODISTANCE (`02_beyond_gm_prep.py, 03_beyond_gm_distances.py`). These beyond-gray-matter distances are quantified in mm units and we included voxels that are maximally 0.7 mm away from any gray matter border. Then, we created a collated distance file, combining the equi-volume depth metric together with the beyond-gray-matter distances (`04_beyond_gm_collate.py, 05_beyond_gm_stitch.py`). This file is used to plot cortical depth profiles together with voxels above and below the gray matter in **Figures 6–7** and **Figures A.1–A.2** (`01_singlesub_depth_vs_T2star.py, 02_singlesub_depth_vs_T1.py, 01_group_depth_vs_T2star.py, 01_group_depth_vs_T1.py`) using Matplotlib (Hunter, 2007).

### 2.6 Flat Maps

We first used LayNii v2.2.0 LN2_MULTILATERATE program to inject a local coordinate system within our segmented regions. The computation of the flat local coordinates involves a combination of computing geodesic distances with respect to four equally distant points lying on a circle of a given diameter centered around our regions of interest and applying a set of linear algebraic equations to a vector field. Further details of the flattening method are outside the scope of the current manuscript and will be elaborated in a future work. However, interested readers can refer to the interim description available at https://thingsonthings.org/ln2_multilaterate.

This flat local coordinate system is abbreviated with U and V letters, which are visible in **Figure 3A**. We then used LN2_PATCH_FLATTEN program to combine the voxel-wise UV coordinates with the equi-volume cortical depth measurements (abbreviated as D coordinate, see **Figure 3B**) to flatten cylindrical (30 mm in diameter) cortical patches within our regions-of-interest (`02_patch_flatten.py`). The end result of this procedure is a full continuous mapping between flat cortex space (UVD) and the original folded cortex space (XYZ) (see **Figure 3C-D**). The UVD coordinates are denoted by the letters “U”, “V”, and “D” based on two reasons. First reason is that the letters “U” and “V” are coming from the computer graphics “UV mapping” term. UV mapping (Botsch et al., 2010) refers to a multitude of methods that find a mapping between a set of points that live in three dimensional coordinates and a set of points that live in two dimensional coordinates. “X”, “Y”, and “Z” are very often reserved for the initial three dimensional space (in our case denoting the initial folded brain coordinates). Therefore, “U” and “V” are selected to denote the two dimensional “flat” coordinates as “X” and “Y” were already taken. Second reason is that we have chosen “D” (as opposed to “W”) because the third coordinate of our flat three dimensional coordinates has a domain specific meaning: “cortical depth”. Therefore, we have extended the “UV mapping” term by adding “D” for depth.

### 2.7 Data and Code Availability

Unprocessed data, sequence parameters (PDFs), and part of the processed data (due to data storage constraints) used in this study are available at: https://doi.org/10.17605/OSF.IO/N5BJ7 under “Data” folder. Processing and analysis scripts used in this study are available at: https://github.com/ofgulban/meso-MRI (v1.0.2 saved at https://zenodo.org/record/7210802).

## 3 Results

### 3.1 Angioarchitectonic information within T_2_* images

Given the high quality of our 0.35 mm T_2_* data, we find that simple inspections reveal clearly visible properties of intracortical vessels. For instance, we are able to visualize penetrating vessel trunks over the cortical surface similar to what is seen with invasive measurement techniques (Duvernoy et al., 1981, Fig. 64). **Figure 4** shows that there are multitude of local T_2_* dips across the cortical surface. While we have manually labeled the most visible dips similar to Duvernoy et al. (1981) in this initial investigation, future studies can improve the detection of such intracortical vessels by developing automatic methods similar to (Bernier et al., 2018; Bollmann et al., 2022; Huck et al., 2019). The unlabeled version of this map for every participant can be found in **Supplementary Figures 5–6** Following Duvernoy’s nomenclature, we call these “intracortical vessels”. These intracortical vessels most likely belong to groups 4 and 5 of the “intracortical veins” based on Duvernoy’s classification (see **Supplementary Figure 9)**. We think that these are likely to be veins because when we overlay the T_2_* images together with the T_1_ contrast (see **Figure 2D-E**), we find many “dark tubular shapes” appearing only in the T_2_* but not in the T_1_ images. In light of the blood motion artifact, it is conceivable that arteries might appear as very dark in T_2_* images too. However, we think that the blood motion is slow enough to not give rise to strong blood motion artifacts from the intracortical arteries. Therefore, we believe that we are predominantly seeing the intracortical veins in the T_2_* images. The reason that we believe the veins belong to group 4 and 5 is that they typically penetrate through the majority of the cortical thickness. The group 4 veins stop at the white matter, while the group 5 veins continue to dive through the white matter. See **Supplementary Figure 9** and compare it with the numbered arrows in **Supplementary Figure 2** where arrows 3 and 8 are putative examples of group 5 veins, and the other arrows are putative examples of group 4.

**Figure 4:**
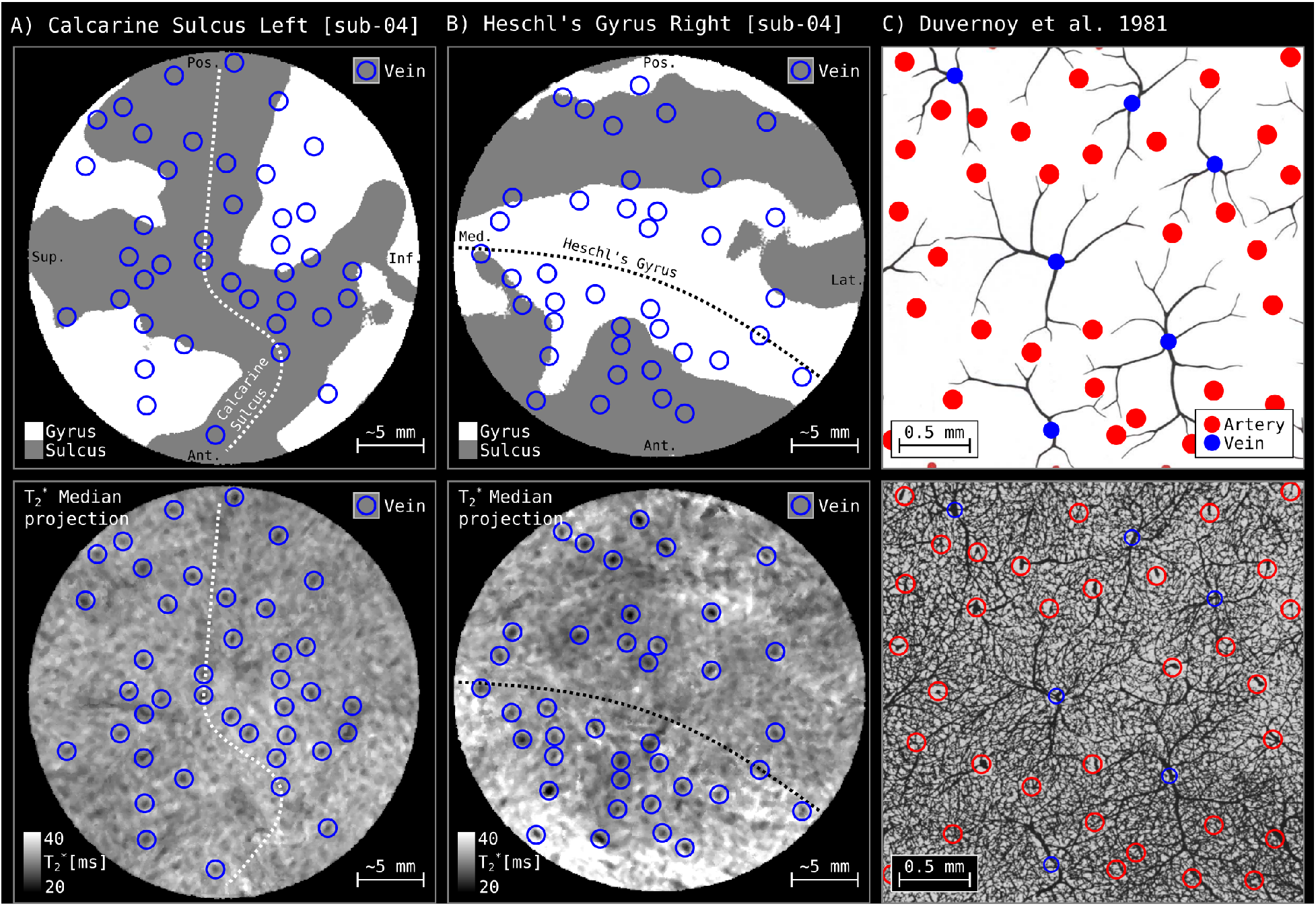
Comparison between in vivo flattened cortical surface representation of T_2_* measurements and ink-gelatin perfusion images from Duvernoy et al. (1981). **(A–B)** Curvature (top) and median-projected T_2_* measurements (bottom) for two example cortical patches. Median projection is performed to summarize all cortical depths into a single image. **(C)** Manually labeled schematic (top) and original ink injection images (bottom) for a sample patch of tissue as reproduced from Duvernoy et al. (1981). Note that there is an order of magnitude difference in the scale bars between A-B and C panels.

Browsing superficial gray matter reveals large T_2_* differences due to the presence of pial vessels that lie above the cortex. **Figure 5A** shows such pial vessel effects sampled from the superficial depth in and around the calcarine sulcus. Branching impressions of the pial vessels can easily be seen as contiguous low T_2_* regions. These dark stripes visible in the flat T_2_* image might be due to the deoxyhemoglobin in veins affecting the T_2_* signal outside of their trunks (Bause et al., 2020), but it could also potentially be due to the blood motion artifact in arteries (as previously discussed). Upon carefully inspecting the MRI signal behavior and the quality of our segmentation and coregistration, we conclude that the dark stripes are likely the result of the former. In addition, **Figure 5B-C** cross-sections reveal the extent of T_2_* darkening can be up to approximately a quarter of the cortical depth. A few large penetrating intracortical vessels are clearly visible in these cross sections as well.

**Figure 5:**
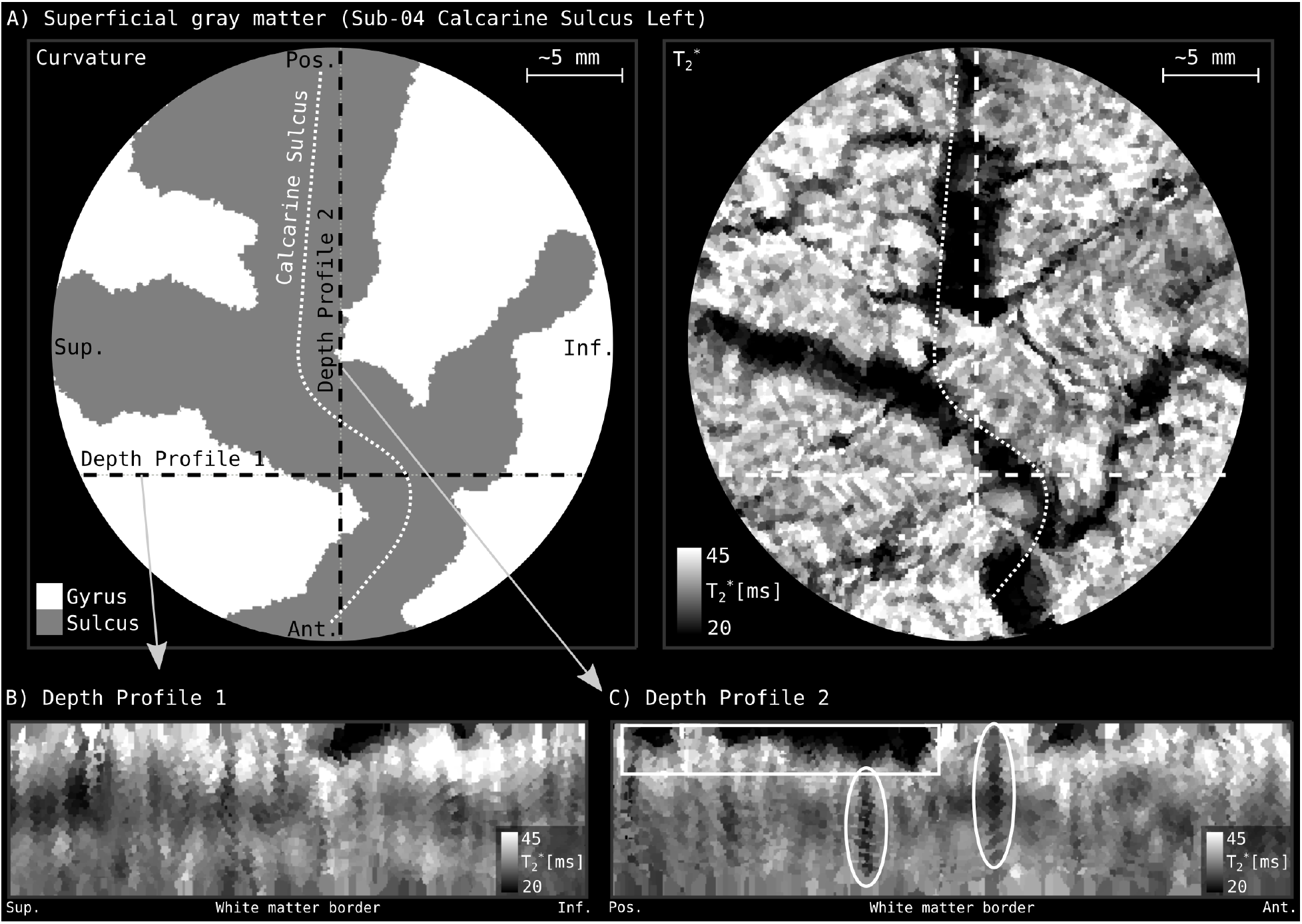
Comparison of flattened T_2_* and T_1_ measurements in a single subject (sub-04). **(A)** Measurements across the cortical surface at the superficial cortical depth. The color associated with each pixel in these images come from exactly one folded brain voxel (processed at 0.175 mm isotropic resolution). T_2_* effects of the pial vessels can be seen as dark impressions in the middle image. **(B–C)**, Measurements for two different cross-sections of cortex. Laminar variations, pial vessel effects (rectangles), and intracortical veins (ellipses) are all visible.

Aside from the very dark T_2_* vascular impressions on the superficial depths and vertical tubes that travel through the cortical thickness, laminar structures are also visible as horizontal bands in **Figure 5B-C**. For instance the stria of Gennari (Fulton, 1937; Gennari, 1782; Glickstein & Rizzolatti, 1984) is visible as T_2_* darkening around the middle of the cortical thickness of the visual cortex. However, we refrain from attributing this laminar structure only to neuronal layers because Duvernoy et al. (1981) and Pfeifer (1940) showed that there are angioarchitectonic layers within the cortex as well.

### 3.2 Cortical measurements of T_2_* at 7 T

Our primary focus was to acquire cortical T_2_* measurements that are valuable for optimizing and interpreting several MRI sequences such as T_2_* -weighted anatomical or functional imaging (Cohen-Adad et al., 2012; Deistung et al., 2013; Markuerkiaga et al., 2021; Marques et al., 2017). For instance, high-resolution fMRI studies make use of the T_2_* values across cortical depths to model unwanted vein effects. Therefore, 0.35 mm iso. resolution of our in vivo quantitative imaging dataset is valuable. In addition, optimal T_2_* -weighting parameters are known to vary across brain regions, and previously reported gray matter T_2_* values are biased towards the occipital cortex (Peters et al., 2007). Therefore, to built a quantitative basis for further optimization and interpretation of the MRI signal, we report T_2_* values across cortical depths for both the visual and auditory areas. We also provide R_2_* variants of our figures in **Supplementary Figure 10–11** for convenience of the reader.

In **Figure 6**, we plotted single-participant T_2_* values across cortical depths for the calcarine sulci and Heschl’s gyri in both hemispheres. It can be seen that there is a T_2_* dip at the middle depth of the gray matter in the calcarine sulci, while a similar dip is not visible for Heschl’s gyri. This observation follows what is visible to the naked eye in **Figure 2E**, namely there is a layer of low T_2_* values within the calcarine sulcus (Barbier et al., 2002; Budde et al., 2011; Duyn et al., 2007; Federau & Gallichan, 2016; Fukunaga et al., 2010; Kemper et al., 2018; Sánchez-Panchuelo et al., 2012; Zwanenburg et al., 2011). This observation holds for every participant (see **Supplementary Figure 2)** and is qualitatively consistent across the left and right hemispheres of each participant. It can be said that average T_2_* is biologically perturbed across cortical depths.

**Figure 6:**
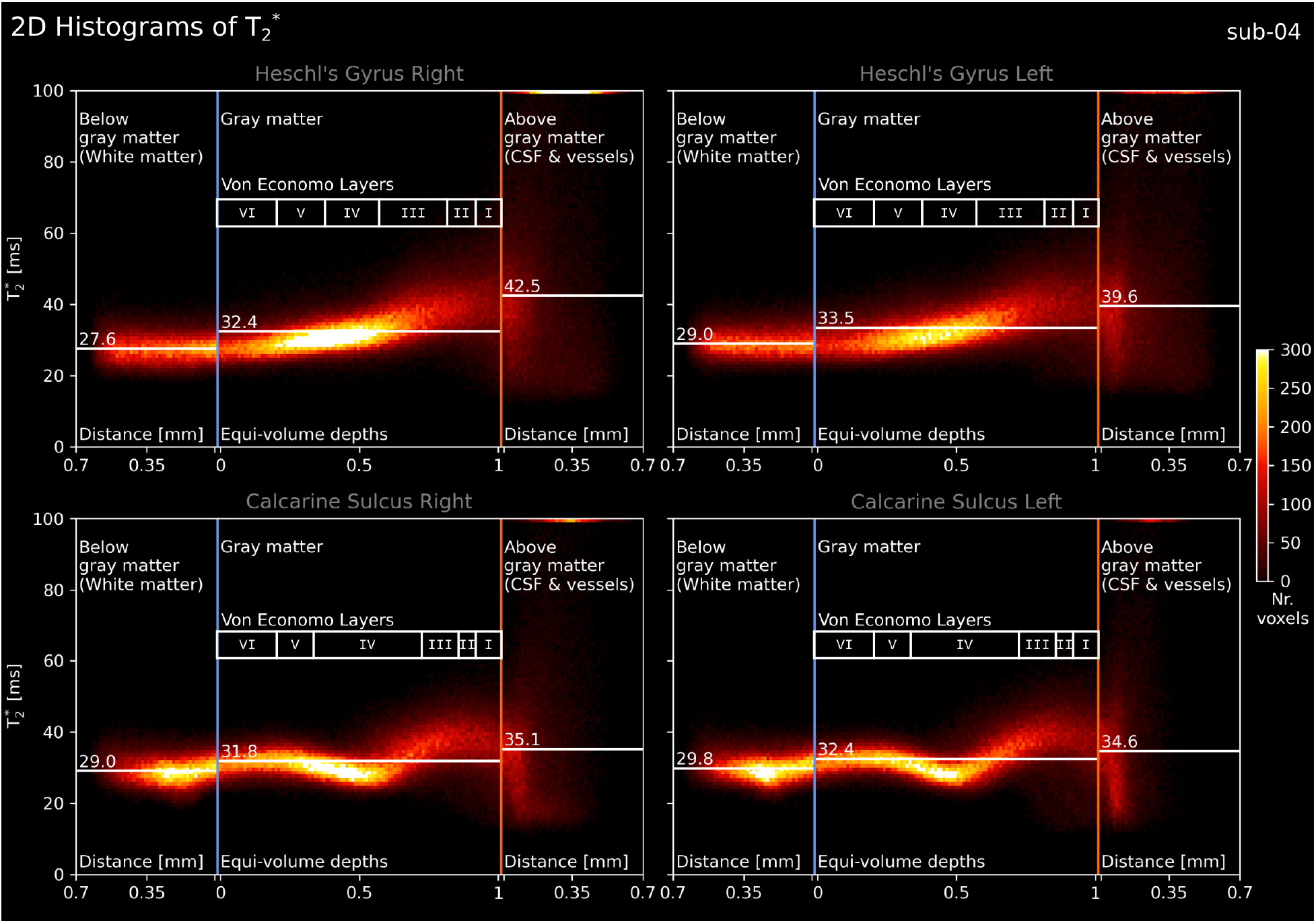
2D histograms of T_2_* as a function of cortical depth for a single participant (sub-04). The vertical lines show gray matter borders: blue for the inner gray-matter border and orange for the outer gray-matter border (see **Figure 2D-E**). Horizontal white lines show the average T_2_* for different tissue sections: below gray matter, gray matter, and above gray matter. Note the topmost bins within the above gray-matter section showing very high T_2_* voxels that are out of range. The “Von Economo Layers” scale is intended as an approximate guide to which cytoarchitectonic layer is most likely being measured rather than clear borders on this individual (Von Economo & Horn, 1930). See **Supplementary Figure 10** for the R_2_* variant. See figure supplements showing each individual T_2_*, R_2_*, and S_0_ at https://osf.io/n5bj7 under *Supplementary Figures* folder.

Looking at the group results in **Figure 7**, we observe that T_2_* variance increases towards the superficial depths in every participant. This could be due to the increased partial voluming with both pial vessels (which causes low T_2_*) and CSF (which causes high T_2_*) that lie above gray matter, and indicates T_2_* is not only neurobiologically biased but also heteroskedastic across depths. Thus, when modelling cortical MRI signals (Havlicek & Uludag, 2020; Markuerkiaga et al., 2021; Uludag et al., 2009), care must be taken not only to account for varying T_2_* but also accounting for the varying amounts of T_2_* variance across cortical depths. In contrast, when considering the white matter voxels below the gray matter (superficial white matter), variance appears to be similar to the deep gray matter voxels. Overall, we observe slightly reduced T_2_* values in white matter compared to gray matter, which could be attributed to the increased myelin content in white matter as well as increased iron content in U-fibers (Kirilina et al., 2020).

**Figure 7:**
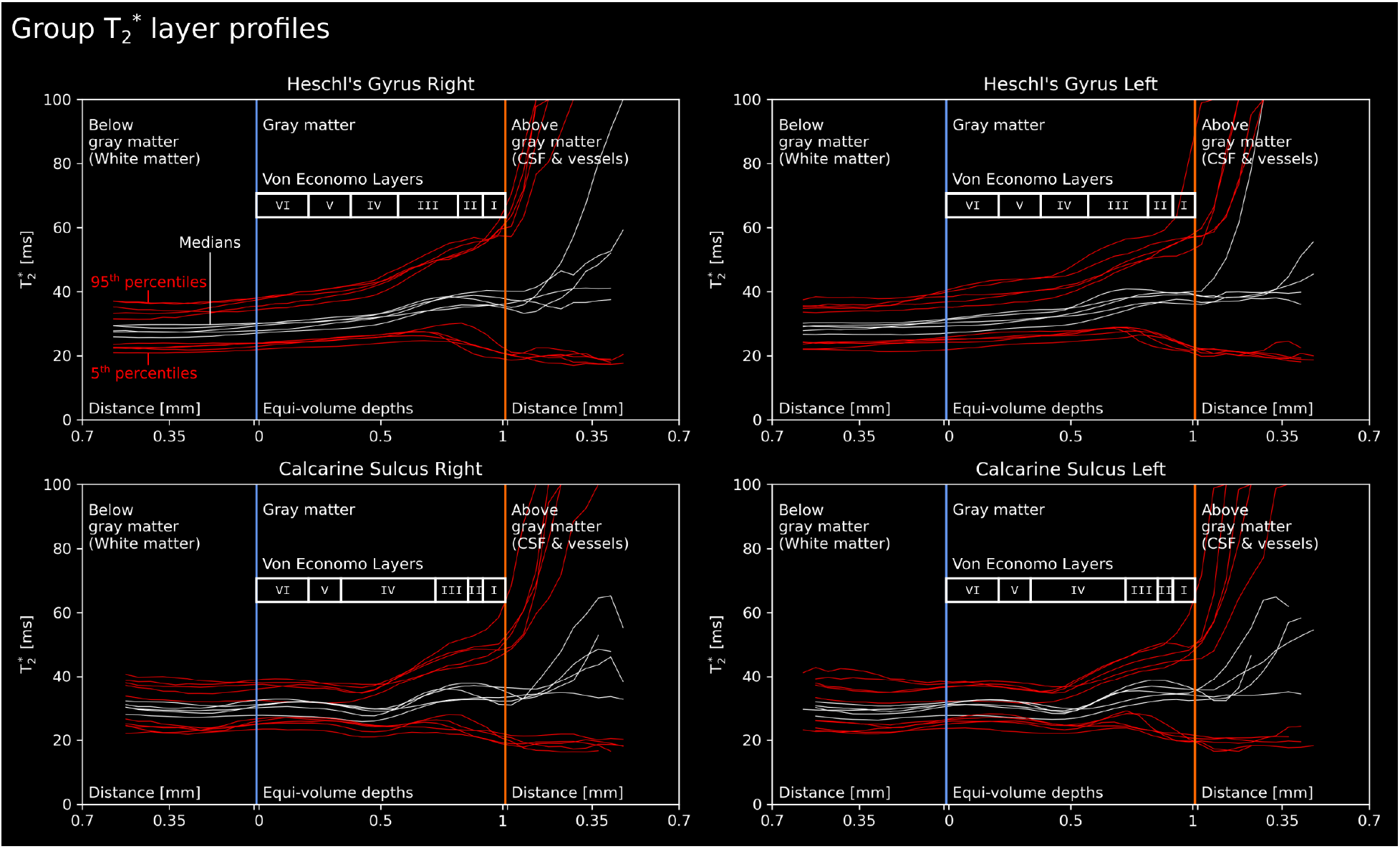
Line plots of T_2_* as a function of cortical depth for all five subjects. Format similar to **Figure 6**. White lines show the median T_2_* calculated for successive bins of cortical depth. Red lines show the 5^th^ & 95^th^ percentiles observed for these bins. See **Supplementary Figure 10** for the R_2_* variant.

## 4 Discussion

In this study, we have demonstrated MRI acquisition and analysis of high-quality mesoscopic anatomical T_2_* data from five living human brains (0.35 × 0.35 × 0.35 mm^3^ resolution). Our freely dataset include quantitative T_2_* measurements from approximately 1/3 of the brain including the primary visual and auditory areas. We provide qualitative demonstrations of fine-scale details captured in our dataset through flat maps. These flat maps reveal clear mesoscopic cortical substructures of the living human brain such as layers and intracortical vessels, and these are discernible without any statistical analysis. We also, for the first time, report quantitative gray matter T_2_* values at 7 Tesla across cortical depths together with superficial white matter, as well as above pial surface measurements for the visual and auditory cortices.

### 4.1 Mesoscopic Properties Observed in Visual and Auditory Cortices

Our mesoscopic T_2_* images reveal clear cortical substructures. Two main types of cortical substructures are visible: layers and vessels. With regards to the cortical layers, there appears to be a substantial difference between the visual and auditory cortices. In the visual cortex, a distinct layer around the middle gray-matter depth is clearly visible as has been previously shown with in vivo MRI at mesoscopic resolutions (Barbier et al., 2002; Budde et al., 2011; Duyn et al., 2007; Federau & Gallichan, 2016; Fukunaga et al., 2010; Kemper et al., 2018; Lüsebrink et al., 2021; Zwanenburg et al., 2011). This structure is the stria of Gennari (Fulton, 1937; Gennari, 1782; Glickstein & Rizzolatti, 1984; Stüber et al., 2014). On the other hand, in the auditory cortex, an equally distinct layering was not immediately visible. This is consistent with post-mortem imaging studies (Wallace et al., 2016; Wallace et al., 2002) and previous sub-millimeter attempts to identify layers within the auditory cortex (De Martino et al., 2015; Dick et al., 2012; Dick et al., 2017; Wasserthal et al., 2014). The difference between the visual cortex layers and the auditory cortex layers in T_2_* and T_1_ images highlight how unique the visual cortex layering is compared to the rest of the cortex. We believe the lack of a clear inflection point in Heschl’s gyrus compared to the calcarine sulcus is real. Thickness variations within Heschl’s gyrus might be suspected to affect the clear appearance of an inflection point, but our equi-volume cortical depth measurements should in theory compensate for such variations. In addition, it is possible that our region of interest included several distinct areas with differing laminar patterns, causing blurring of different inflection points. However, careful visual inspection of the data indicates that a laminar pattern as strongly visible as in the calcarine sulcus is simply not present in our data. Nonetheless, as argued within Kuehn et al., 2017; Skeide et al., 2018; Wallace et al., 2016, mesoscopic in vivo imaging may still be insufficient to capture subtle changes in myeloarchitecture beyond what can be found in the primary visual cortex where layer IV is extremely thick. Future studies might explore improving signal-to-noise in existing measurements (e.g. Lohmann et al., 2010) or increasing spatial resolution even further.

Our mesoscopic T_2_* images also show clear presence of vascular structures within the cortex. This is interesting because seminal neuroscience works highlight laminar cytoarchitecture, myeloarchitecture, and fiber architecture (Turner, 2013a, 2013b), but rarely mention laminar angioarchitecture. It has been shown that the angioarchitecture of the cortex exhibits a laminar arrangement both across depths and over the cortical surface (Duvernoy et al., 1981; Pfeifer, 1940). Specifically within Duvernoy’s work, four angioarchitectural layers are postulated (see **Supplementary Figure 9)**. When the first two angioarchitectonic layers combined, it can be seen that these layers almost equally divide the human cortex. Given that T_2_* contrast has a substantial contribution from the presence of blood, the laminar patterns we are observing within the visual cortex (see **Figure 2E** and **Supplementary Figure 2)** may also reflect the laminar angioarchitecture (also see Koopmans et al. (2011) Fig.6). Indeed, Pfeifer (1940) showed increased density of blood vessels in middle cortical layers, which might be the origin of the T_2_* dip we have observed. More broadly, what constitutes the living brain might be biased by the post-mortem focus within the last century where angioarchitectonic layering within the cortex is partially neglected due to the draining of blood and degradation of blood vessels during fixation and embedding processes. We invite others to make use of our mesoscopic dataset to further study cortical angioarchitecture.

### 4.2 Challenges That Remain for Mesoscopic Imaging

There are several remaining challenges for in vivo mesoscopic MRI. The most obvious challenge is head motion (Jezzard & Clare, 1999; Zaitsev et al., 2015). In our study, all participants were highly experienced for being scanned and were able to keep their heads relatively still within the data acquisition periods. In addition, we used 3D acquisition sequences rather than 2D sequences; in 3D sequences, within-run head motion is less sensitive to motion-induced spin-history effects. Potential future application of motion correction methods can also mitigate such motion effects (Cordero-Grande et al., 2020). Another advantage of 3D sequences is that they generally enable higher resolution and signal-to-noise ratio (Poser et al., 2010; Stirnberg et al., 2017). As a result of using 3D acquisitions and experienced participants, we achieved highly detailed images of cortical substructures, even those as small as intracortical vessels. This result shows that mesoscopic MRI is practical when head motion is minimized and when using 3D acquisition sequences with sufficient signal-to-noise ratio, participant preselection. However, it would be critical to use additional strategies to minimize head motion of less experienced participants, such as prospective motion correction (Lüsebrink et al., 2017; Maclaren et al., 2012; Mattern et al., 2018; Stucht et al., 2015; Zaitsev et al., 2017), 3D fat-based motion navigators (Federau & Gallichan, 2016; Gallichan et al., 2016; Gretsch et al., 2020), 3D printed head casts (Power et al., 2019), and field monitoring (Barmet et al., 2008; Eschelbach et al., 2019). We think that more advanced head motion minimization methods have potential to yield crisper mesoscopic images and broaden the application of mesoscopic MRI to a larger audience. While we fully acknowledge that we performed retrospective head motion correction rather than prospective motion correction (see **Supplementary Table 1)**, we believe that our results make a good argument for the importance of participant selection when prospective motion correction methods are inaccessible (see **Supplementary Figure 2. 3, 4)**. We have consistently collected high quality data by combining pre-selection of experienced participants with shorter “fMRI-style” (maximum 10-15 min.) acquisition durations that allow for resting in-between runs.

In our study, we used conservative amounts of image acquisition acceleration. This resulted in somewhat lengthy but still feasible anatomical acquisition durations (10–15 min). The choice of minimal acceleration was deliberate in order to minimize artifacts, maximize signal-to-noise ratio, and thereby establish a benchmark dataset. However, a persistent challenge for mesoscopic imaging is long data acquisition times. Faster acquisitions leveraging e.g. echo planar imaging (Sanchez Panchuelo et al., 2021b) are especially needed for clinical populations. From a methods development perspective, our densely sampled high-resolution multi-echo dataset might be useful for efforts in which a dataset is subsampled and methods are developed to optimally recover the full dataset from limited data, such as parallel imaging or model based reconstructions (Ye, 2019).

Even in an ideal case of no head motion, in vivo MRI presents a unique challenge, namely, effects due to the motion of blood. Especially within large arteries, blood reaches very high speeds (Rahman Rasyada & Azhim, 2018). Such moving blood is known to cause imaging artifacts when the time between the phase-encoding and readout (frequency encoding) stages of the MRI signal acquisition gets longer (Larson et al., 1990; Wehrli, 1990). An obvious case where blood motion artifacts are visible is multi-echo acquisition, where each phase-encoded line of k-space is measured using several successive readouts to acquire multiple echos. Our ME GRE images have captured this artifact in unprecedented detail (**Supplementary Figure 1)**. Importantly, this artifact is not specific to our anatomical multi-echo acquisition. Any MRI sequence that has long (> 4 ms) time difference between the phase-encoding and readout stages is bound to have this artifact, given that the arterial signal is not effectively nulled. Especially in echo planar imaging (EPI) readouts (Bernstein et al., 2004) commonly used for fMRI, where the image is acquired within very long readout durations (e.g. > 30 ms), this blood motion artifact will strongly exhibit itself. Fortunately, the blood motion artifact is limited to regions near large arteries (> 1 mm diameter). However, studies that rely on accurate and precise quantification of the brain tissue (e.g. see literature within Mancini et al., 2020), such as multi-parametric mapping and in vivo histology (Edwards et al., 2018) should be aware of this artifact. In addition, studies that rely on weighted contrasts, such as T_2_* -weighted, should also be cautious of this artifact because the arterial signal can be smeared towards the gray matter, causing artifactually hyperintense regions. Moreover, the artifact may corrupt contrast locally by introducing dark regions in the actual position of an artery as its signal is displaced away from its trunk. This can cause arteries to appear similar to the venous signal in T_2_* -weighted images. Thus, dark regions in T_2_* -weighted images - especially near the pial surface - cannot be conclusively labeled as veins, as has been suggested (Kay et al., 2019; Moerel et al., 2018; Olman et al., 2007). In order to dissociate whether the dark T_2_* voxel are arteries or veins, we recommend tracking the trunk of the vessels to determine which cerebral artery or large collecting vein they are connecting to.

### 4.3 Future Neuroscience Applications for Mesoscopic MRI

Our study suggests exciting possibilities for mesoscopic imaging. From a clinical perspective, mesoscopic in vivo imaging of human cortical architecture creates the opportunity to develop a novel set of biomarkers for the healthy development of adult cortical laminar structure and its disruption in neurological and psychiatric disorders (McColgan et al., 2021). Prime candidate conditions for mesoscopic imaging include well-established but subtle abnormalities in laminar structure that currently lack clear imaging correlates such as layer-specific neuronal loss in amyotrophic lateral sclerosis (Braak et al., 2017) and Huntington’s disease (Rüb et al., 2016), as well as developmental disruption of cortical layering in focal cortical dysplasia (Blümcke et al., 2011). From a cortical parcellation perspective, future analyses may investigate whether inter-regional variability in laminar structure can be leveraged for parcellating cortical areas using approaches like those used for cytoarchitectonic data (Dinse et al., 2015; Gulban et al., 2020; Morosan et al., 2001; Schleicher et al., 1999; Zachlod et al., 2020). With additional acquisition time, it is clear that mesoscopic coverage of 1/3 of the brain can be easily increased to include the whole brain by scanning different slabs and stitching them together. From an fMRI perspective, imaging the angioarchitectonic substructure of the cortex is necessary for building biophysical models of the fMRI signal (Akbari et al., 2022; Báez-Yánez et al., 2020; Cheng et al., 2019; Havlicek & Uludag, 2020; Polimeni & Lewis, 2021). The spatial variability of fMRI signals due to mesoscopic angioarchitecture is not well understood. The effect of intracortical veins has been the focus of fMRI signal modelling so far (Havlicek & Uludag, 2020; Heinzle et al., 2016; Markuerkiaga et al., 2016; Markuerkiaga et al., 2021; Uludag et al., 2009), although empirical studies have shown considerable contributions from pial veins (Bause et al., 2020; Kurzawski et al., 2022; Olman et al., 2007; Turner, 2002; Winawer et al., 2010). Further, several sub-millimeter fMRI studies have found heterogeneous response profiles across cortical depths (Chen et al., 2013; Fracasso et al., 2018; Kashyap et al., 2018). Thus, the angioarchitecture information in our mesoscopic images might help extend classical models of fMRI signals to consider the heterogeneity of the underlying vasculature and improve the interpretability of laminar BOLD fMRI profiles.

## 5 Appendix A

### 5.1 MP2RAGE T_1_ cortical depth profiles

We highlight that our primary focus for acquiring the T_1_ contrast was for cortical gray matter segmentation. Therefore, we have used a speed-optimized MP2RAGE sequence that provided good contrasts for cortical segmentation purposes. However, we believe that there could be some value in reporting our findings. T_1_ contrast is clearly shown to be related to myelination (Leuze et al., 2017; Morawski et al., 2018) and therefore used for delineating areal borders (Cohen-Adad et al., 2012; Deistung et al., 2013; Dick et al., 2012; Dinse et al., 2015; Geyer et al., 2011; Haast et al., 2016; Marques et al., 2017; Sanchez Panchuelo et al., 2021b; Sánchez-Panchuelo et al., 2012).

We plot single-subject T_1_ measurements across cortical depths for the calcarine sulci and Heschl’s gyri in both hemispheres in **Figure A.1**. We observe a dip of T_1_ values in middle of the cortical thickness in the calcarine sulci, but a similar dip is not visible for Heschl’s gyri. This T_1_ dip is harder to identify in individual brain images compared to the dip in T_2_*. Looking at the variance of T_1_ across cortical depths in group results **Figure A.2**, we find that variance does not increase as drastically as in T_2_* profiles (see **Figure 7)**. This suggests that T_1_ is less sensitive to laminar architecture than T_2_*. Looking at the above gray-matter voxels, it can be seen that the variance increases, but less than in the T_2_* profiles. This is because vessels show on average similar T_1_ to gray matter, while CSF has a higher T_1_ (Zhang et al., 2013). We also provide R_1_ variants of our figures in **Supplementary Figure 12–13** for convenience of the readers.

A comparison between T_1_ and T_2_* images leads to interesting observations (e.g. **Fig.2D-E**). CSF, arteries, and veins can be considered as the three main tissue types that lie above gray matter. When looking at the T_2_* images, vessels can be seen as regions with low T_2_* while CSF has very high T_2_*. In **Figure 2D-E**, looking at the low T_2_* voxels in T_1_ images reveal that large (> 1 mm diameter) arteries and veins seem to have very different T_1_ values. This difference might be due to the higher velocity of the blood in the arteries compared to veins affecting proper inversion of the arterial signal during MP2RAGE acquisition. However, aside from the arteries, it can be seen that most of the low T_2_* voxels above gray matter are not visible in MP2RAGE T_1_ images. This observation is interesting because it shows that T_2_* contrast, which is sensitive to macromolecular content and iron, is highly advantageous for localizing veins compared to T_1_ contrast, which is mostly sensitive to the macromolecular content.

Another interesting yet subtle detail is the consistent asymmetry between the average gray matter T_1_ values across hemispheres. It can be seen that the right hemisphere average gray matter T_1_ is higher in all participants compared to their left hemisphere when comparing Heschl’s gyri. Such asymmetry is not visible when comparing the calcarine sulci. Given the calcarine sulci are much closer in space compared to Heschl’s gyri, we attribute this average gray matter T_1_ difference to residual the transmit field inhomogeneities based on Marques and Gruetter (2013).

**Figure A. 1:**
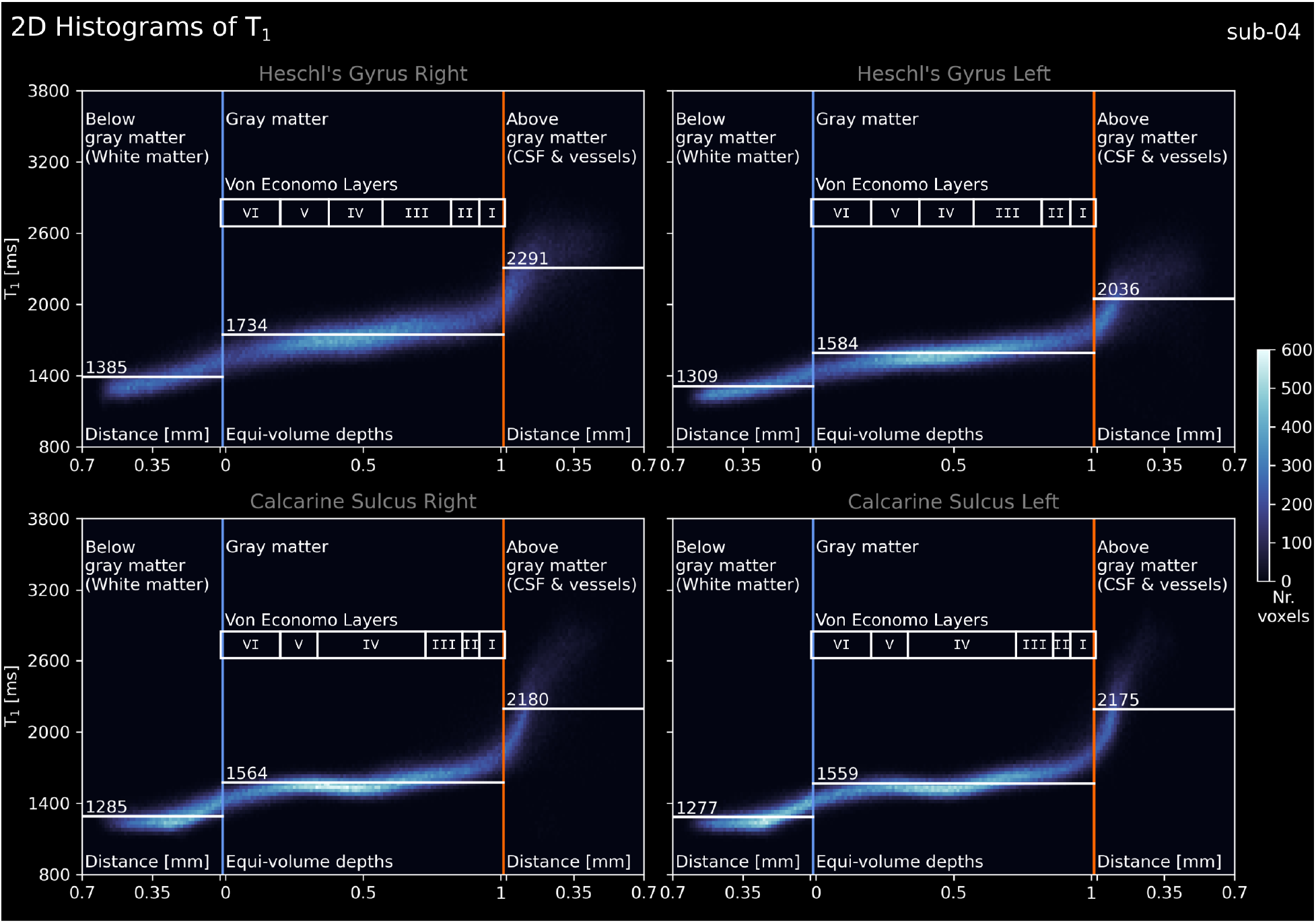
2D histograms of T_1_ as a function of cortical depth for a single subject (sub-04). The vertical lines show the gray-matter borders: blue for the inner gray-matter border and orange for the outer gray-matter border (see **Figure 2D-E**). Horizontal white lines show the median T_1_ for different tissues: below gray matter, gray matter, and above gray matter. The “Von Economo Layers” scale is intended as an approximate guide to which cytoarchitectonic layer is most likely being measured rather than clear borders on this individual (Von Economo & Horn, 1930). See figure supplements showing each individual T_1_, R_1_, and S_0_ at https://osf.io/n5bj7 under *Supplementary Figures* folder. See **Supplementary Figure 12** for the R_1_ variant.

**Figure A. 2:**
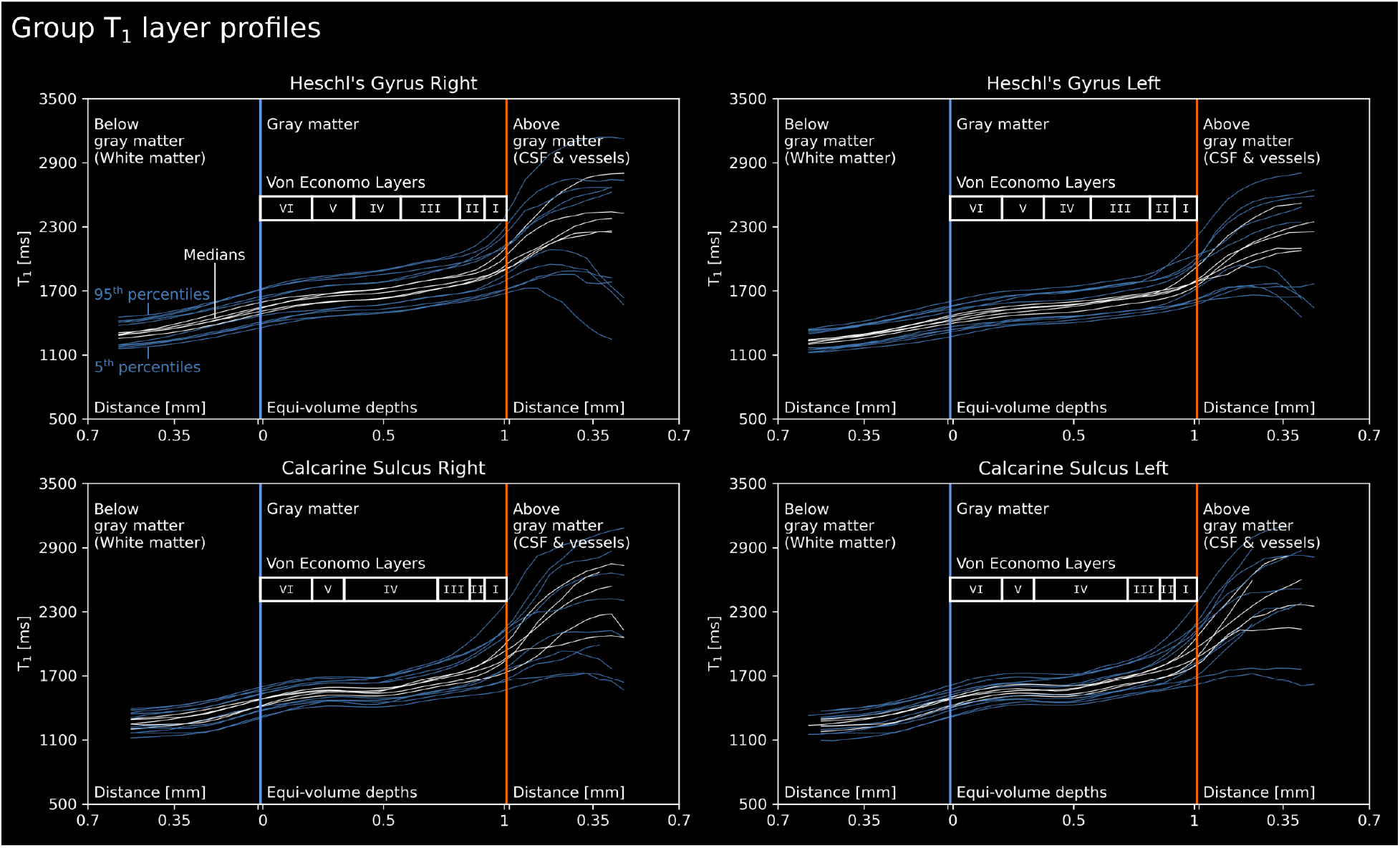
Line plots of T_1_ as a function of cortical depth for all five subjects. Format similar to **Fig.A.1**. White lines show the median T_1_ calculated for successive bins of cortical depth. Blue lines show 5^th^ & 95^th^ percentiles observed for these bins. See **Supplementary Figure 13** for the R_1_ variant.

## Author Contributions

According to the CRediT (Contributor Roles Taxonomy) system.

**Conceptualization:** O.F.G., D.I.

**Data curation:** O.F.G., D.I.

**Formal Analysis:** O.F.G.

**Funding acquisition:** D.I., R.G.

**Investigation:** O.F.G., D.I.

**Methodology:** O.F.G., D.I., K.K., S.B., R.H.

**Project administration:** O.F.G., D.I., K.K.

**Resources:** O.F.G., R.G., R.H., B.A.P.

**Software:** O.F.G., R.H.

**Supervision:** O.F.G., D.I., K.K.

**Validation:** O.F.G., D.I., K.W., S.B.

**Visualization:** O.F.G., K.K., D.I.

**Writing – original draft:** O.F.G., D.I., K.K.

**Writing – review & editing:** O.F.G., D.I., K.K., S.B., K.W., R.H., B.A.P., R.G.

## Acknowledgements

We thank our anonymous reviewers for their helpful comments. We thank Nikola Stikov, Jonathan Polimeni, Jeff Duyn, David Norris, and Robert Turner for their comments on the first version of our preprint. We have benefited from these valuable comments to improve our manuscript. We thank Federico De Martino for his contributions in the early stages of this study. We thank Valentin Kemper, Martin Havlicek, Kamil Uludag, Nikolaus Weiskopf, Logan Dowdle, Jonathan Winawer, Simon Robinson, Korbinian Eckstein, Michelle Moerel for fruitful discussions and their support at various stages of this project. Additionally, we thank Elizabeth DuPre, Gilles De Hollander, Taylor Salo, and Agah Karakuzu for their help with Brain Imaging Data Structure (BIDS) organization of our dataset (https://neurostars.org/t/multi-echo-anatomical-mri-bids-questions/17157).

SB was funded from the NHMRC-NIH BRAIN Initiative Collaborative Research Grant APP1117020, and NIH grant 1R01MH111419. R.H. was funded from the NWO VENI project 016.Veni.198.032. K.W. was funded by the Wellcome Trust (215901/Z/19/Z). B.P. was partially funded by the NWO VIDI grant 16.Vidi.178.052, by the National Institute for Health grant R01MH/111444 (PI: David Feinberg) and by the H2020 FET-Open AROMA grant agreement no. 88587. R.G. received funding from the European Union’s Horizon 2020 Framework Programme for Research and Innovation under the Specific Grant Agreement No. 945539 (Human Brain Project SGA3). O.F.G. and R.G. have financial interest tied to Brain Innovation. Scanning was supported by FPN (faculty of psychology and neuroscience) via the MBIC grant scheme. Scanning was performed at the facilities of Scannexus B.V. (Maastricht, Netherlands). T_2_* -mapping was done with an ASPIRE sequence, kindly provided by Simon Robinson via SIEMENS C2P.

## Supplementary Material

**Supplementary Figure 1:**
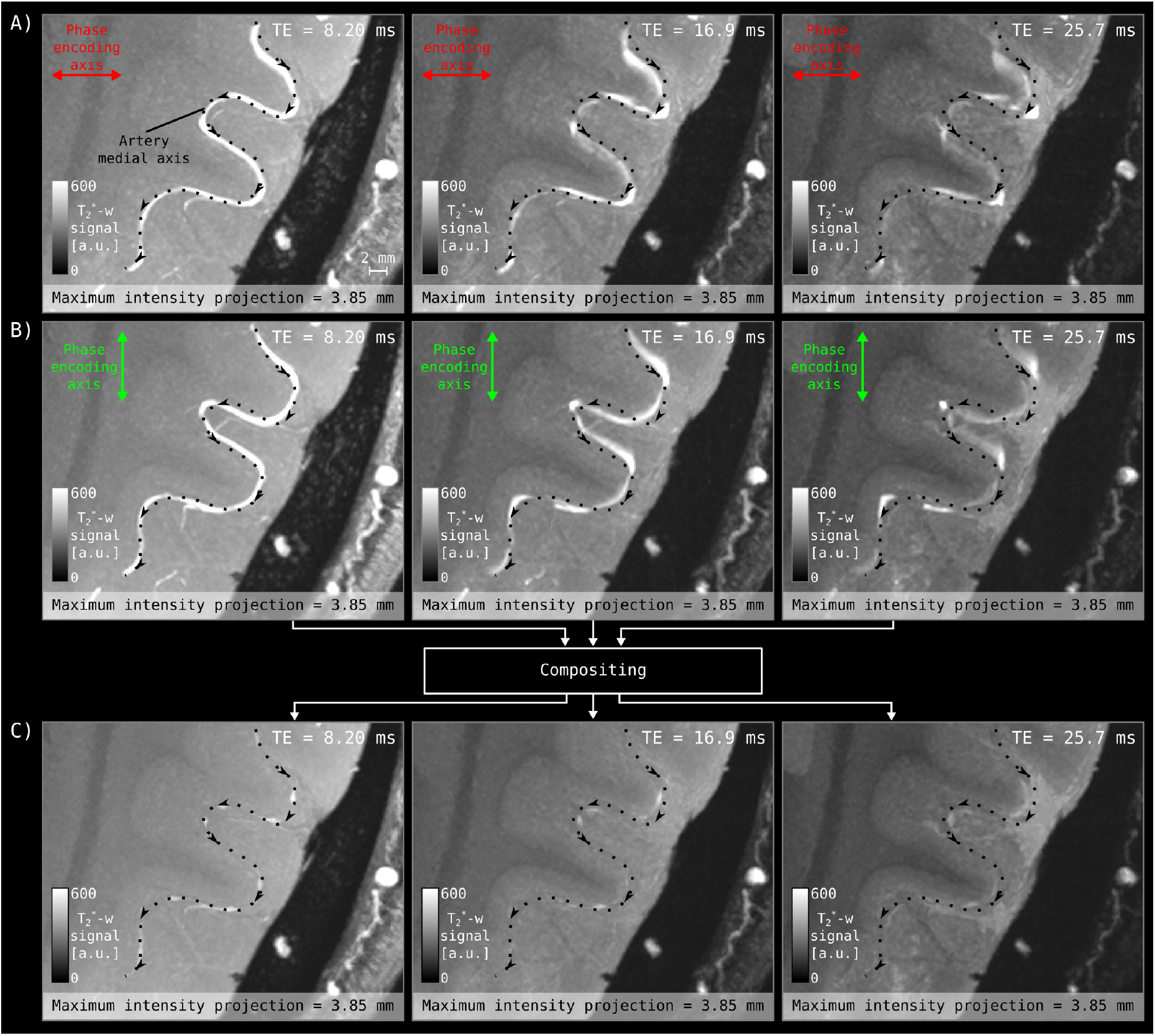
Compositing method to mitigate blood motion artifacts in T_2_* -weighted images. Here we show results from an example participant (sub-03). **(A)** Zoomed-in view of three of the six echos acquired using the right-left phase encoding axis. **(B)** Same echos acquired using anterior-posterior phase encoding axis. See that the blood motion artifact appears along the vector component of the blood flow along the readout axis (which is perpendicular to the phase-encoding axis). **(C)** Result of compositing the acquired images using the minimum operator. We show maximum intensity projection over a small slab (see the first panel for the scale bar) in order to enhance the visibility of the arterial signal. The dotted line superimposed on all images indicates the medial axis of an artery that was manually identified from the shortest TE (3.83 ms) image where the blood motion artifact is minimal. It can be seen that the gray speckles within the dark regions in the Panel A and B are substantially diminished in the Panel C, showing that the compositing improved image signal-to-noise ratio while keeping the tissue boundaries sharp. However, we emphasize that effectiveness of our mitigation method relies heavily on good coregistration across acquisitions.

**Supplementary Figure 2:**
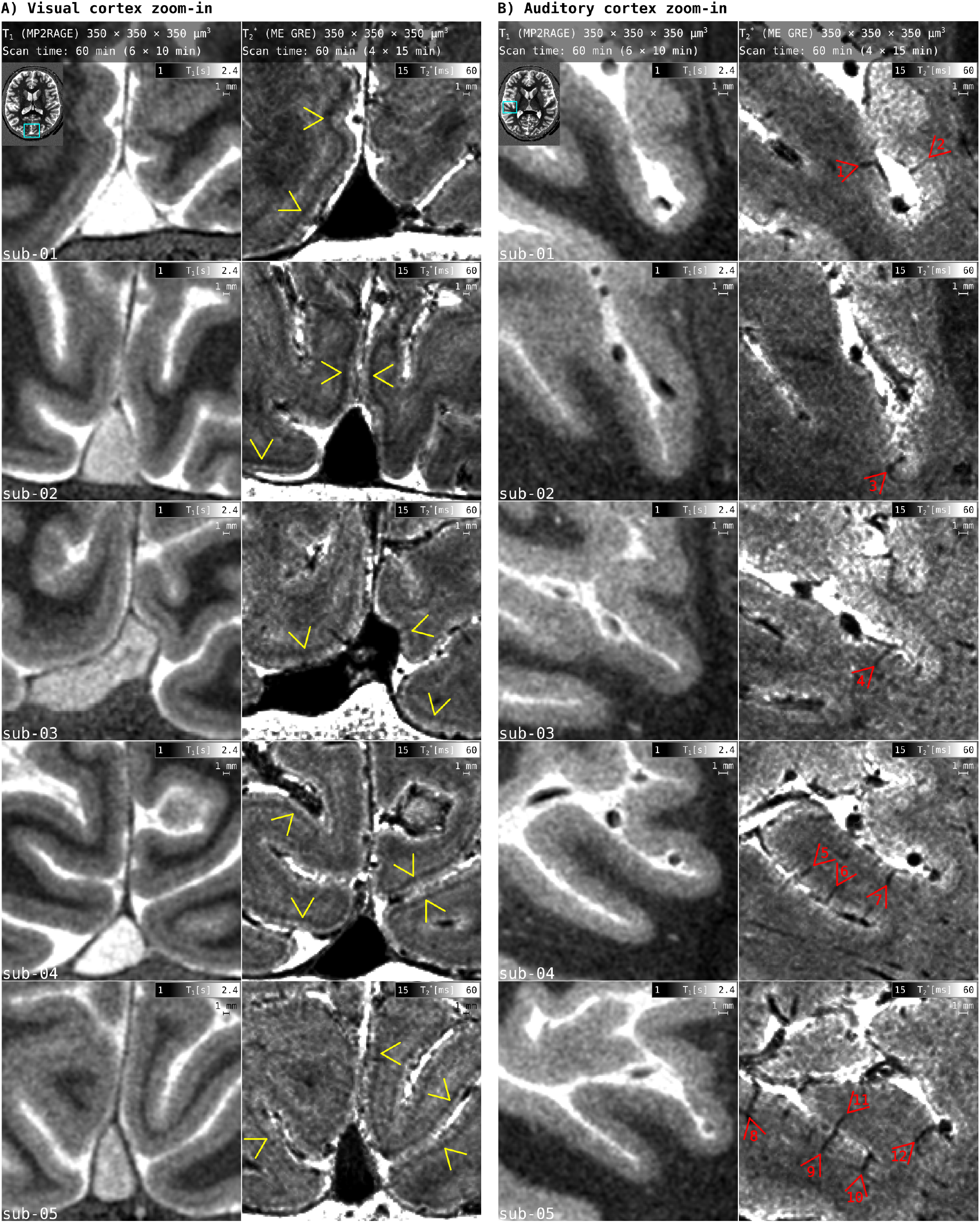
Data quality after preprocessing. Yellow arrows point towards the stria of Gennari. Red arrows point towards intracortical vessels. Arrows 1 & 4 highlight cases in which part of a vessel lies on top of the gray matter. Arrows 9 & 11 highlight cases in which a common vein collects blood from two opposite gray matter chunks. Arrow 12 highlights a case where a smaller vessel connects to a large vein lying at the bottom of a sulcus.

**Supplementary Figure 3:**
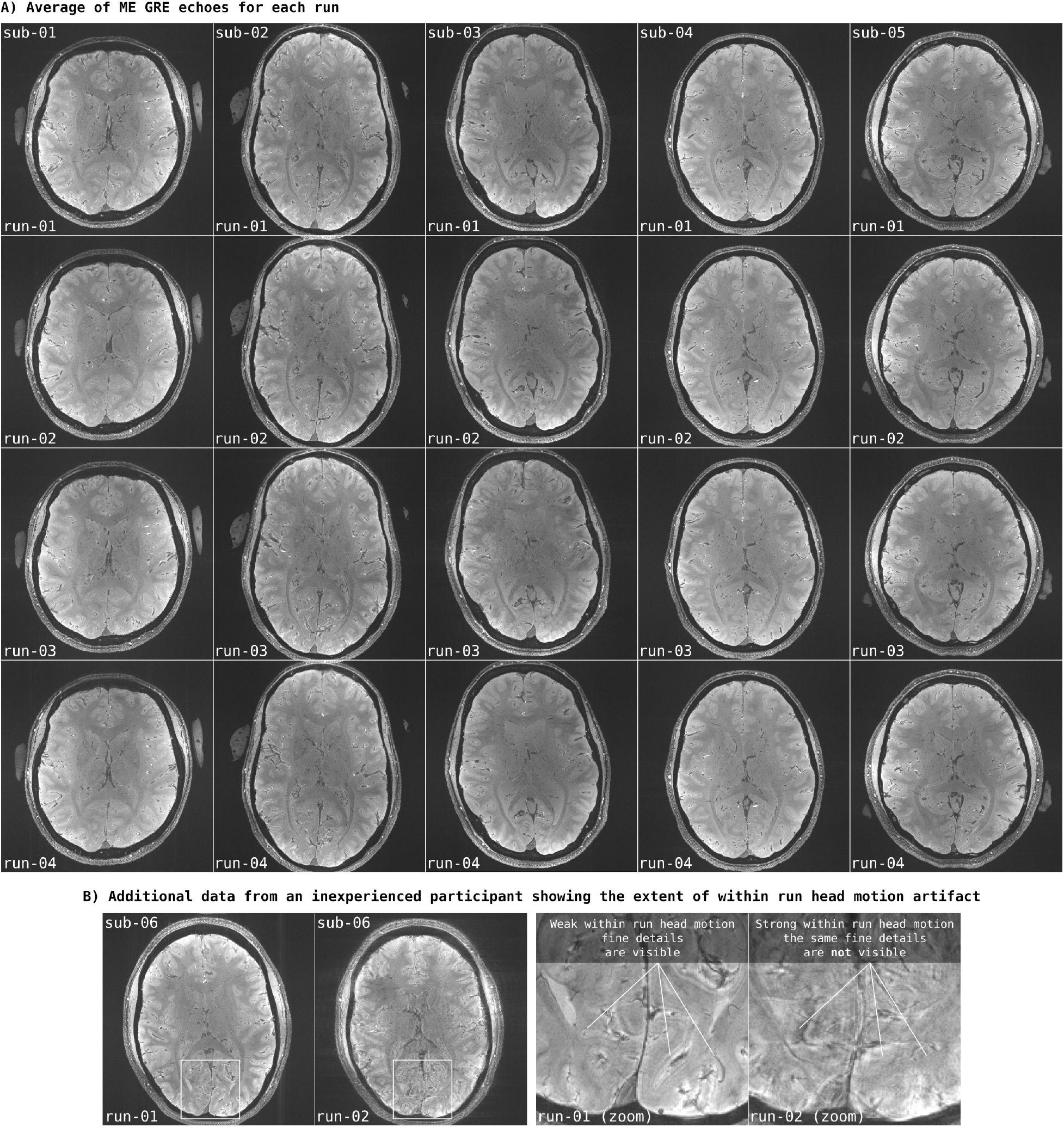
ME GRE data quality for each run before preprocessing. (A) Here we show the average of all echoes at the middle slice of each acquisition. Given the high quality of these data, we ended up discarding no individual runs. (B) Additional data acquired from an inexperienced participant (who did not complete the scanning sessions) to demonstrate within run head motion artifacts (we did not process or analyze this data further). When a participant moves too much during data acquisition, the resulting images have specific types of blur artifacts. Typical approaches to mitigate head motion include prospective motion correction and/or preselecting participants who are able to keep their head still. In the present study, we have employed participant preselection in combination with “fMRI-style” 10-15 minute long single data acquisition run.

**Supplementary Figure 4:**
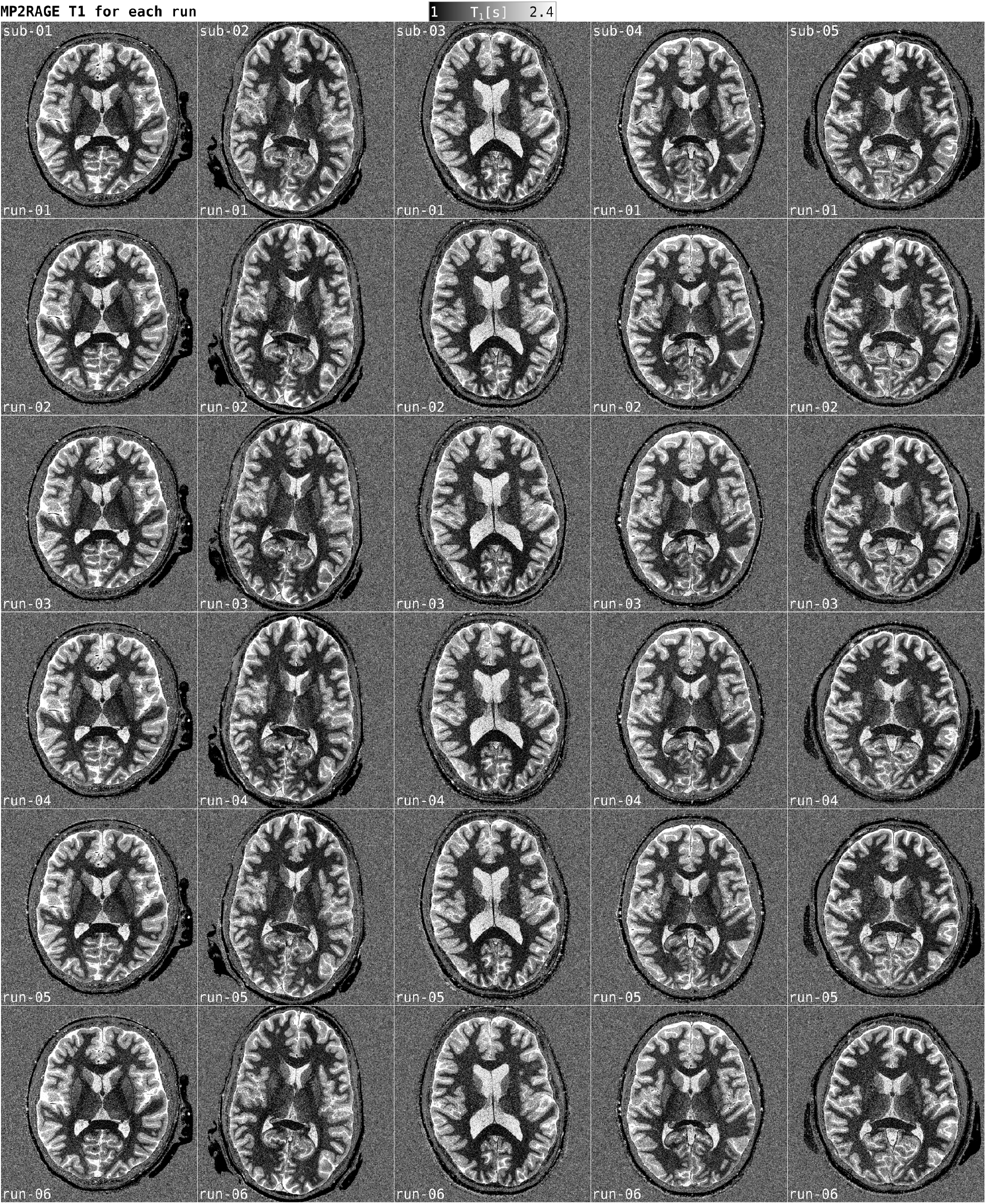
MP2RAGE data quality for each run before preprocessing. Here we show MP2RAGE T_1_ images at the middle slice of each acquisition. The same T_1_ range shown in the colorbar on top is used for all panels. Given the consistent and high quality of these data, we ended up discarding no individual runs.

**Supplementary Figure 5:**
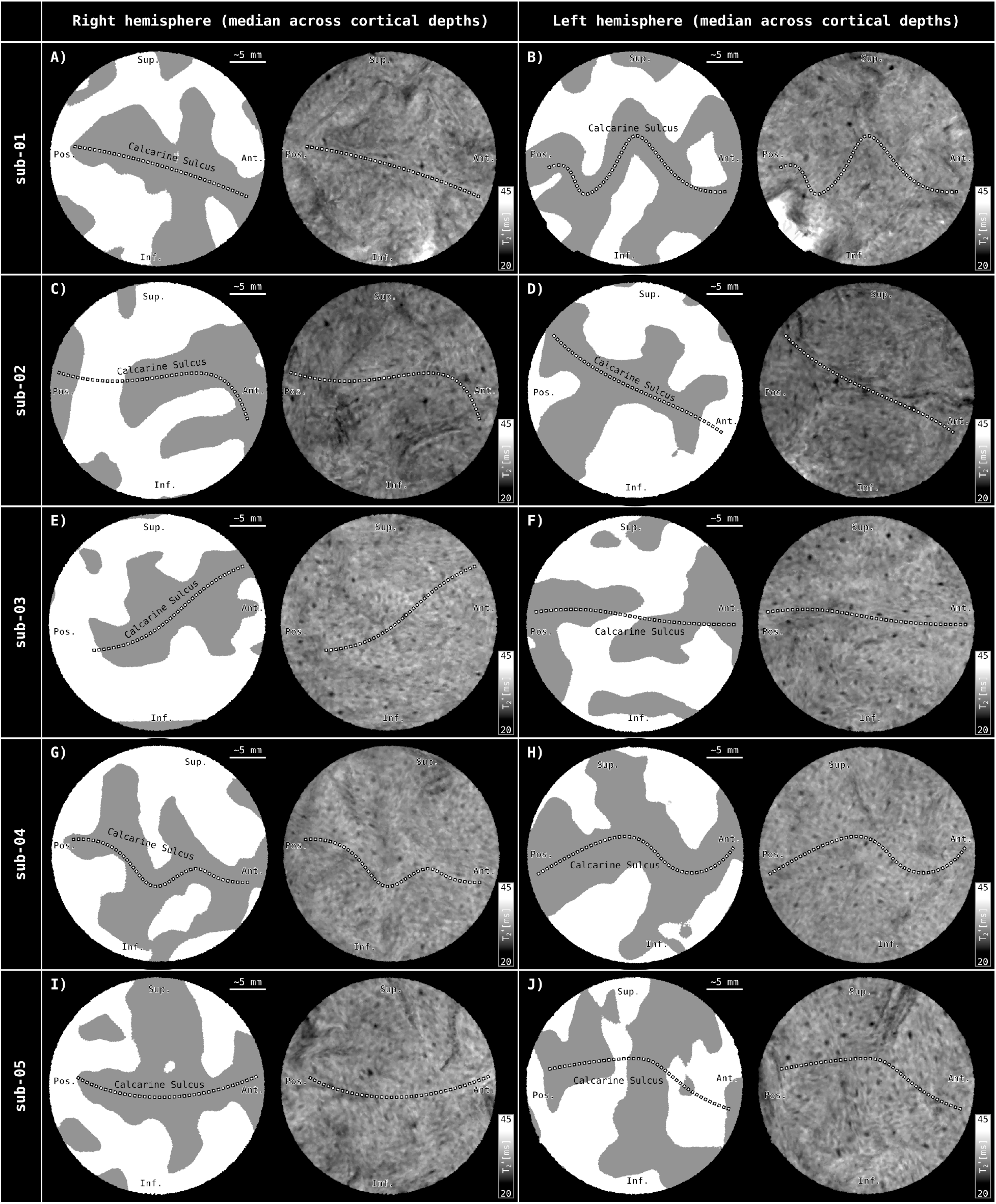
Flattened calcarine sulci of all participants showing median projection across critical depths. Format similar to **Figure 4**.

**Supplementary Figure 6:**
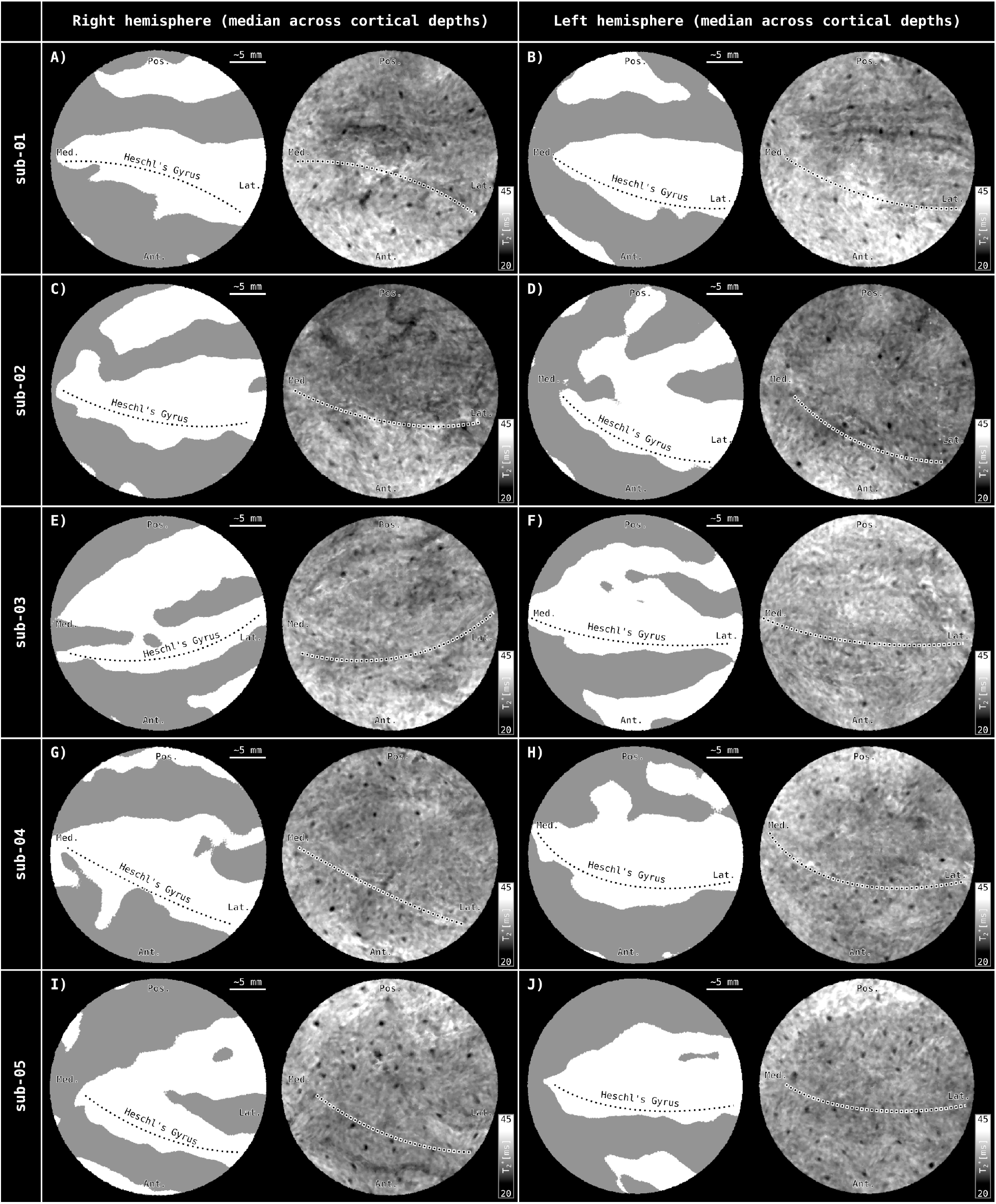
Flattened Heschl’s gyri of all participants showing median projection across critical depths. Format similar to **Figure 4**.

**Supplementary Figure 7:**
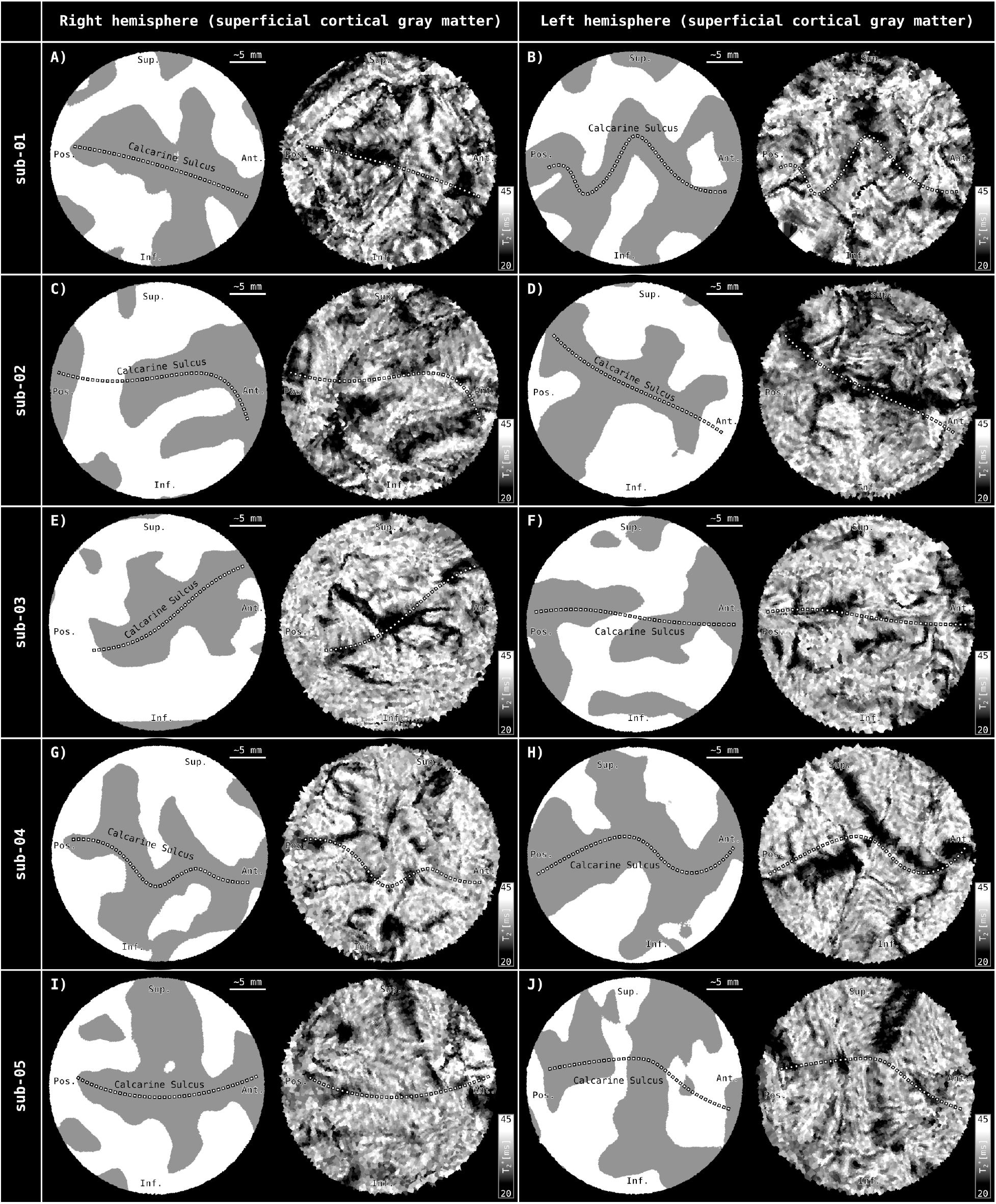
Flattened superficial cortical depths of calcarine sulcus across all participants. Format similar to **Figure 5**.

**Supplementary Figure 8:**
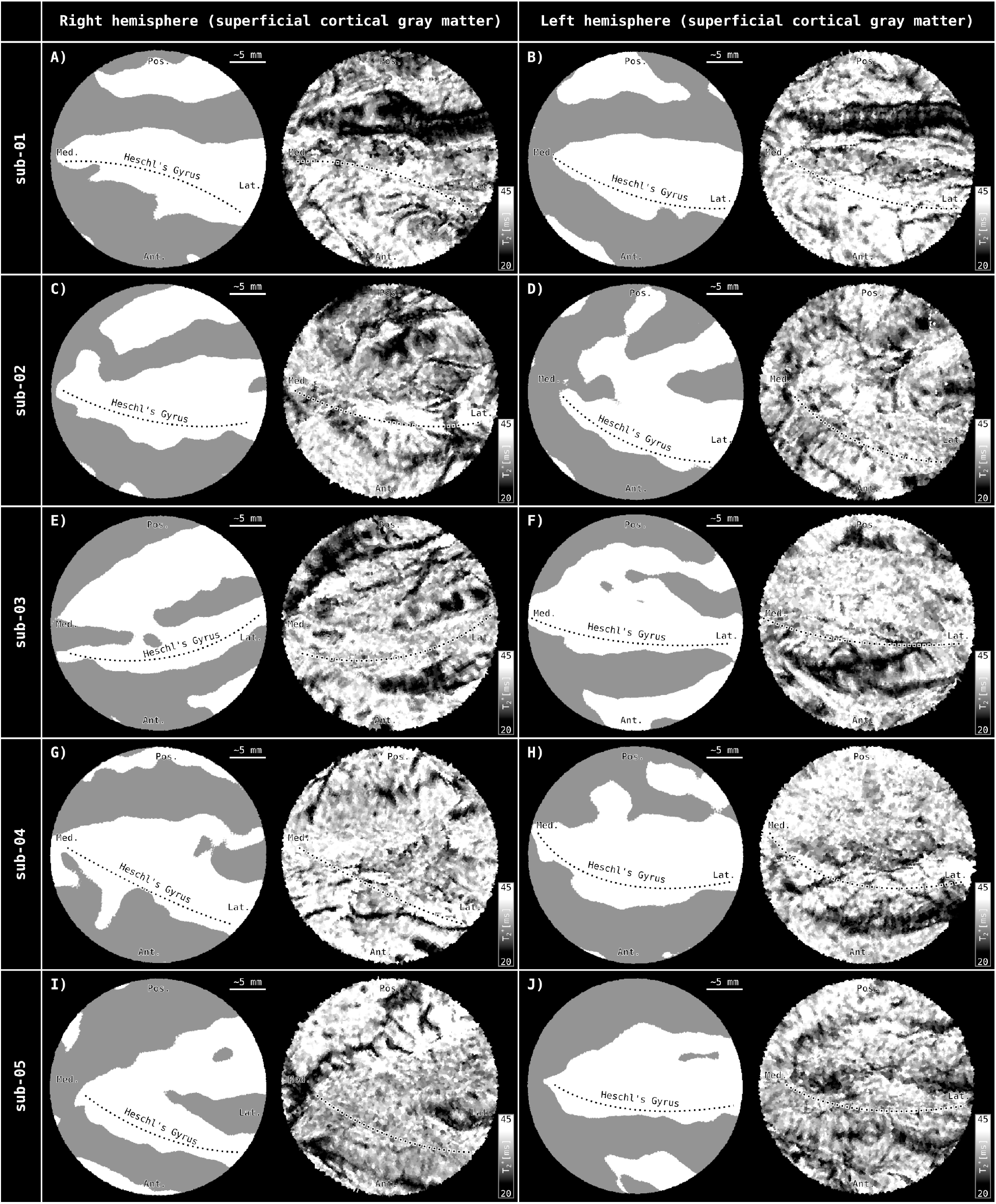
Flattened superficial cortical depths of Heschl’s gyrus across all participants. Format similar to **Figure 5**.

**Supplementary Figure 9:**
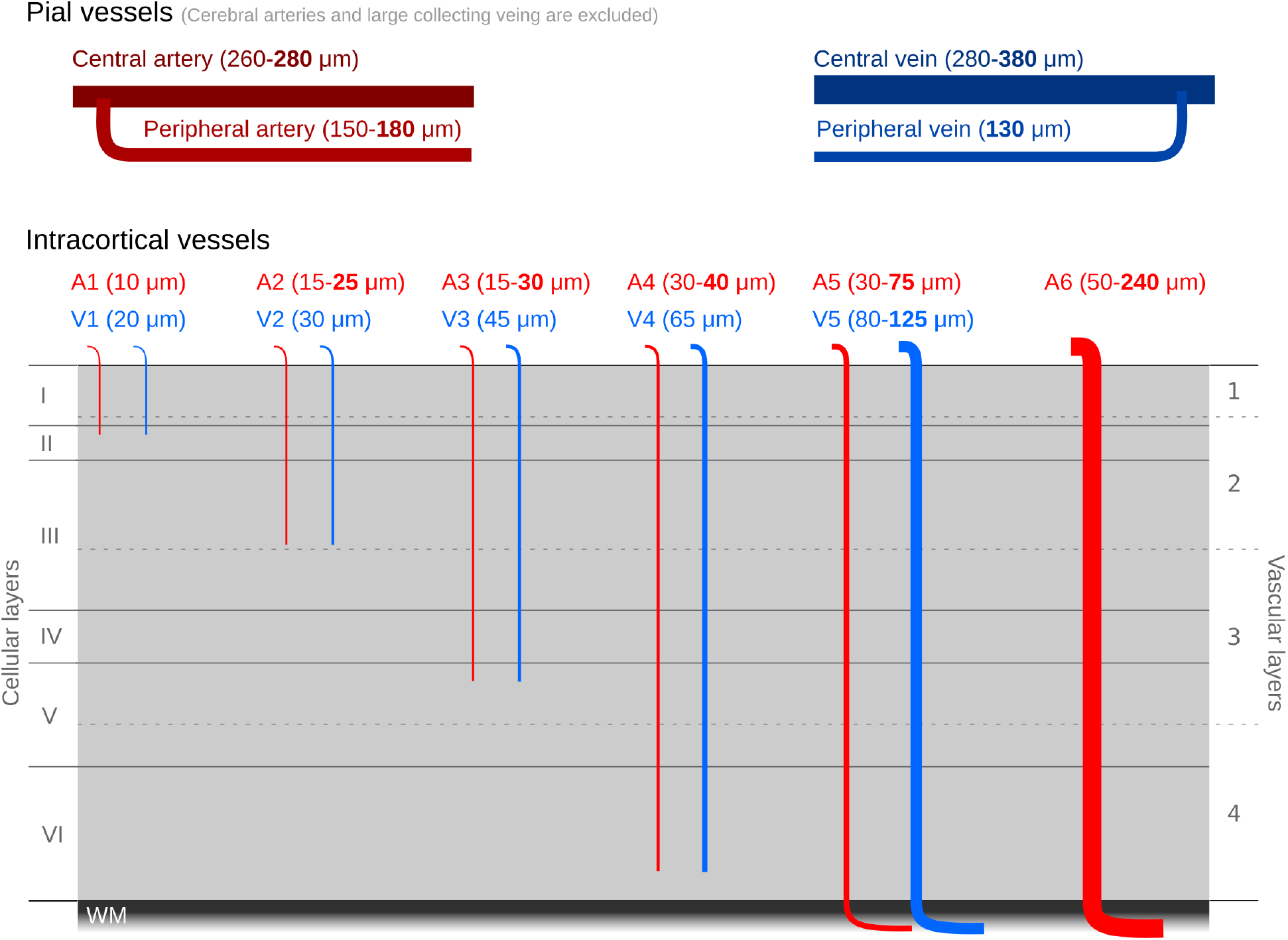
Schematic representation of cortical vessel trunks based on Duvernoy et al., 1981. Information within this figure is taken from the the following sections of Duvernoy et al.: “Cortical Artery Diameters”, ”Superficial Cortical Vein Diameters”, “Intracortical Artery Diameters”, “Intracortical Vein Diameters”, Figure 25, and Figure 52. The vessel diameters reported here are known to be imprecise measurements. As Duvernoy states, (I) *“The diameter of cortical arteries is difficult to evaluate; the results of measurements are debatable and are a function of the pressure of injection and modifications caused by the fixation. Even in vivo … there are large variations in the diameter of cortical vessels according to physiological conditions.”*; and (II) *“Fixation and embedding often greatly deform veins, due to the thinness of their walls; thus, diameter measurements taken after India ink injection are of little value and are often lower than those obtained with vascular casts.”*.

**Supplementary Figure 10:**
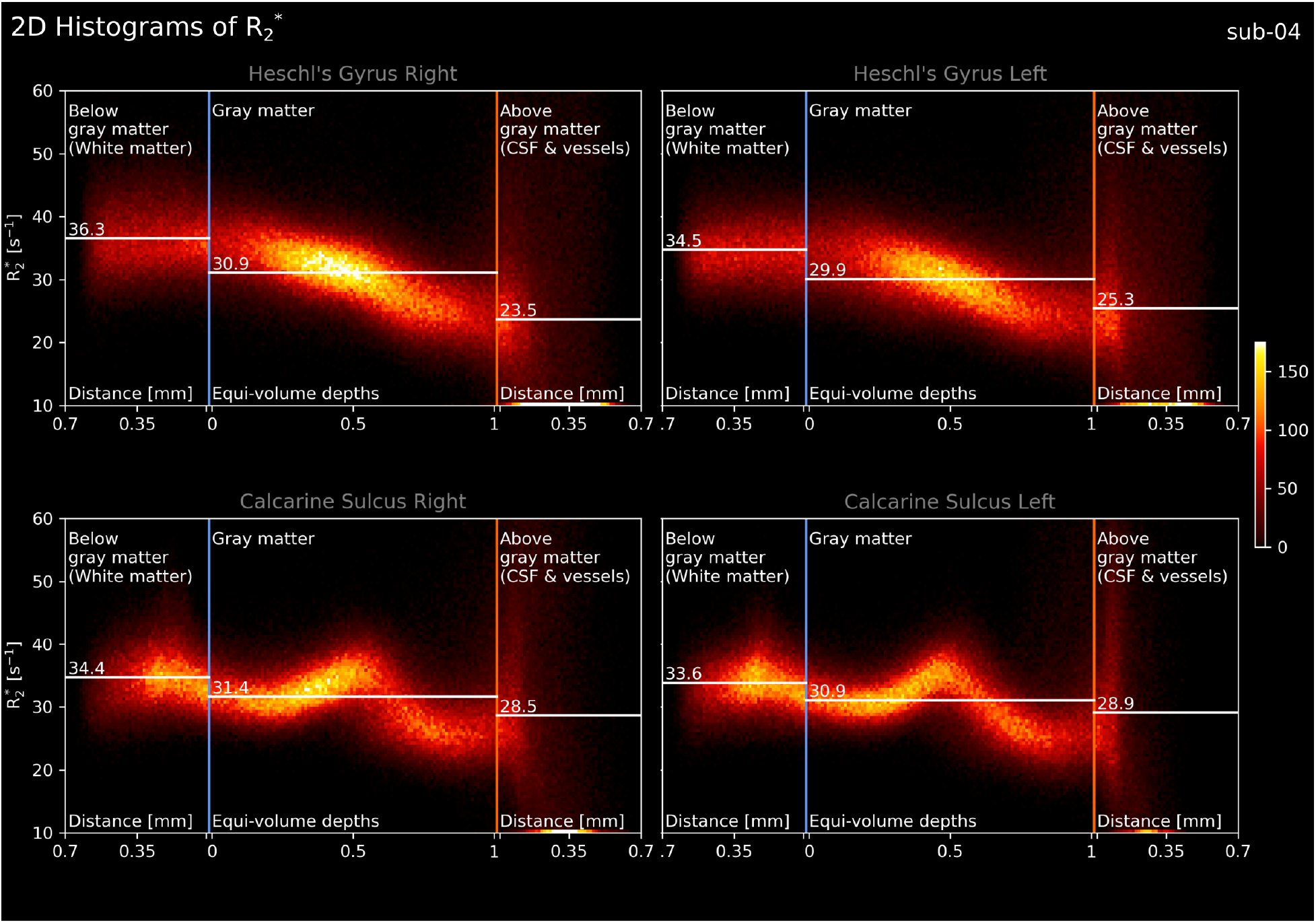
2D histograms of R_2_* as a function of cortical depth for a single subject (sub-04). Variant of **Figure 6**.

**Supplementary Figure 11:**
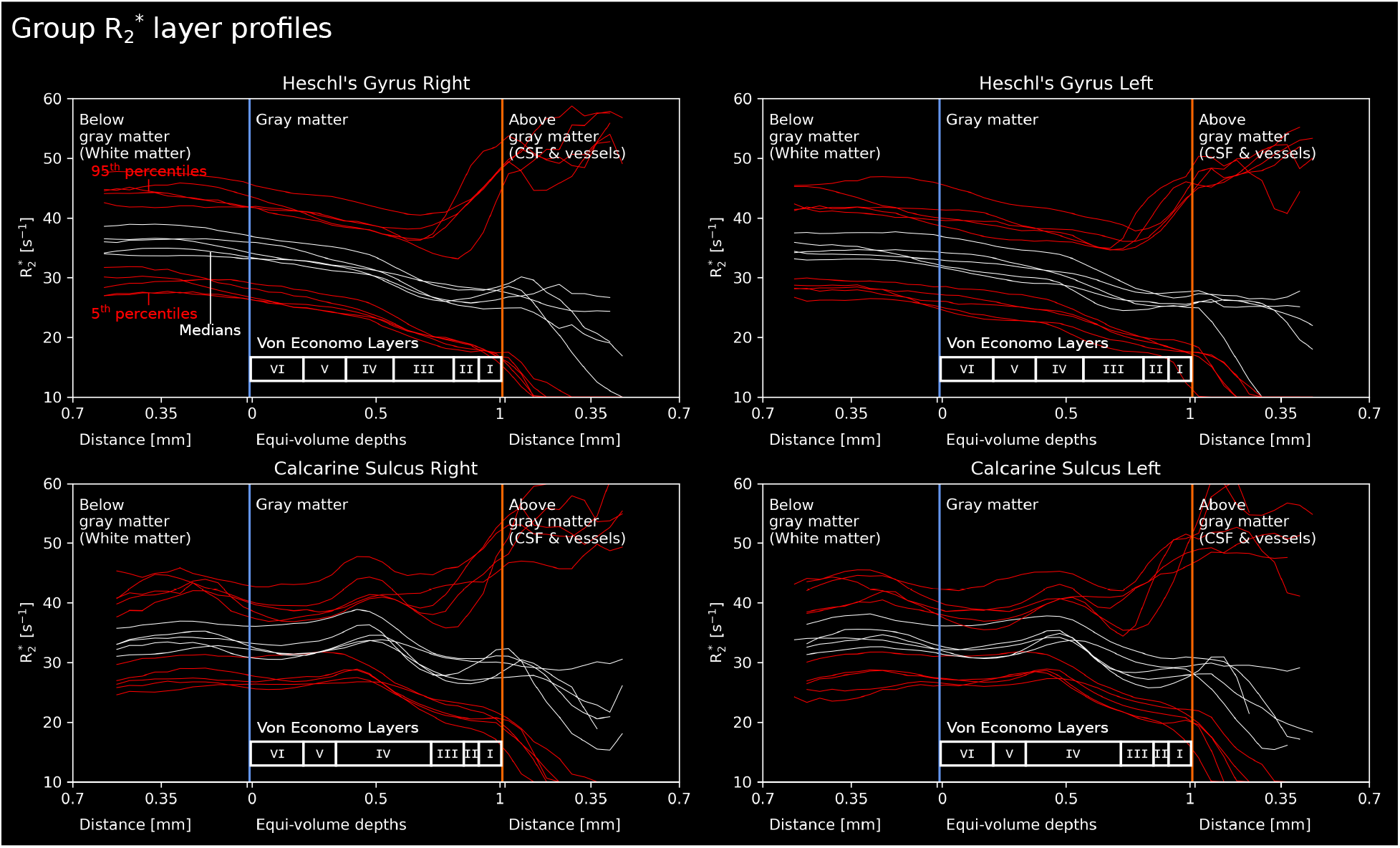
Line plots of R_2_* as a function of cortical depth for all five subjects. Variant of **Figure 7**.

**Supplementary Figure 12:**
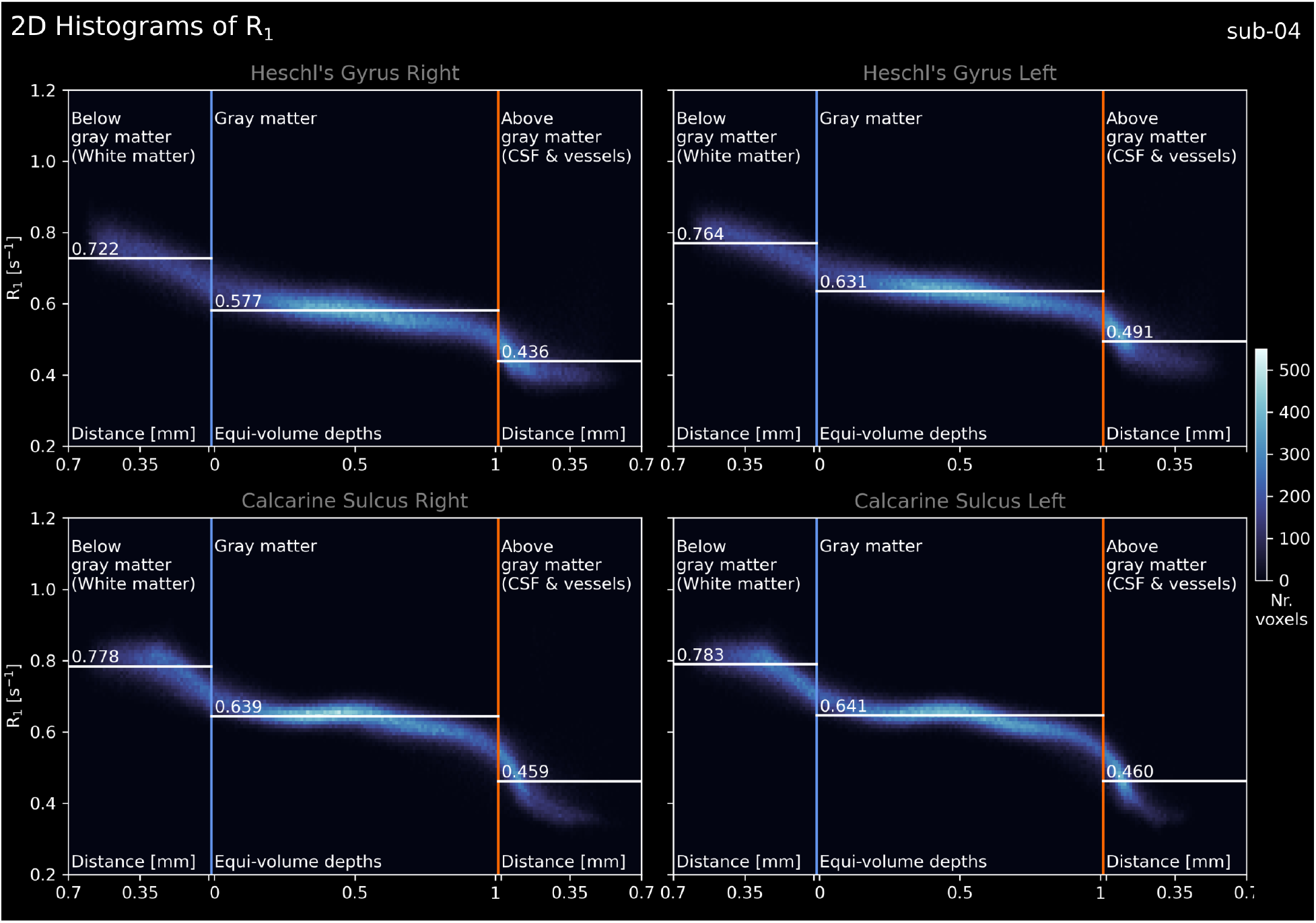
2D histograms of R_1_ as a function of cortical depth for a single subject (sub-04). Variant of **Figure A. 1**.

**Supplementary Figure 13:**
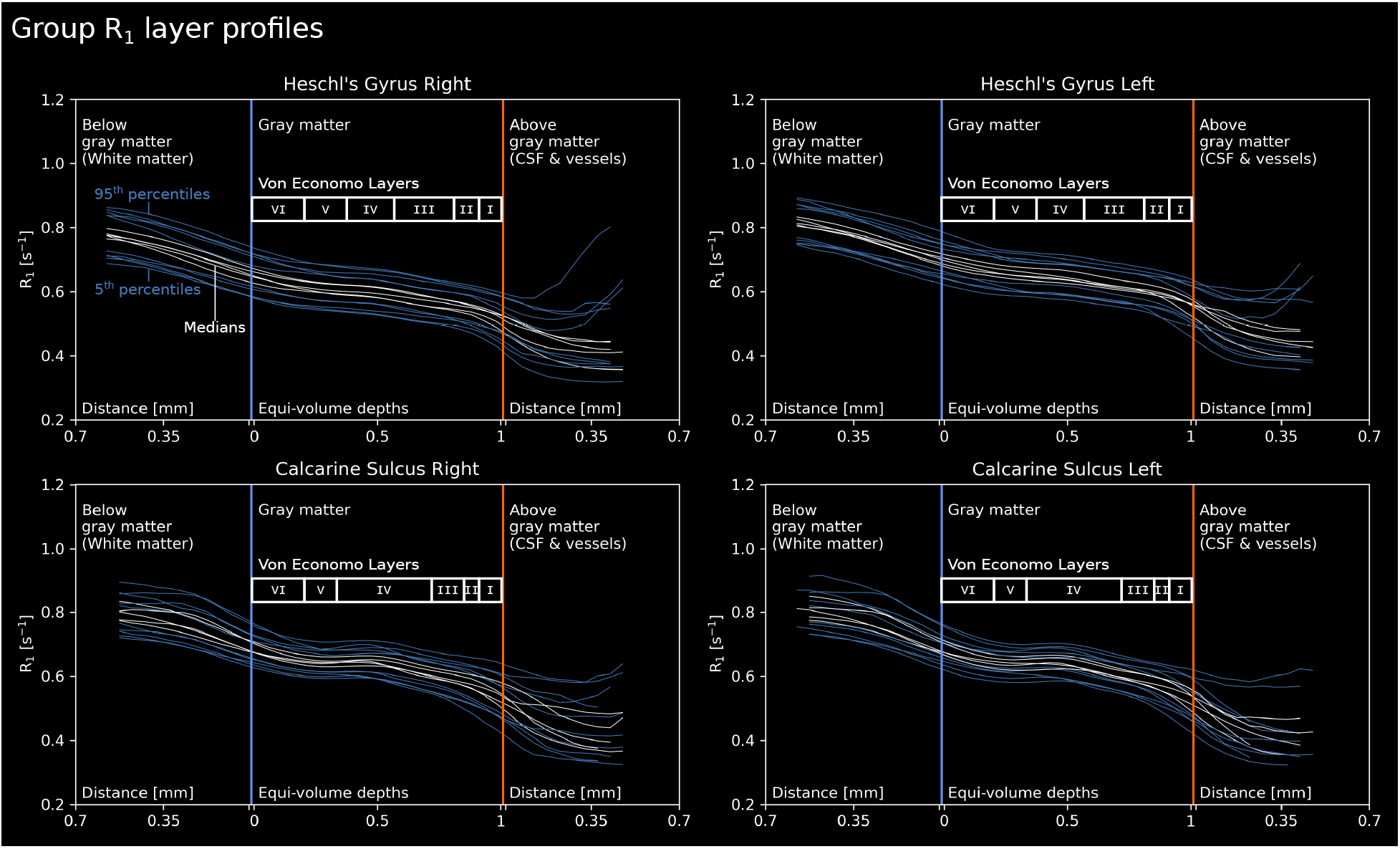
Line plots of R_1_ as a function of cortical depth for all five subjects. Variant of **Figure A. 2**.

**Supplementary Figure 14:**
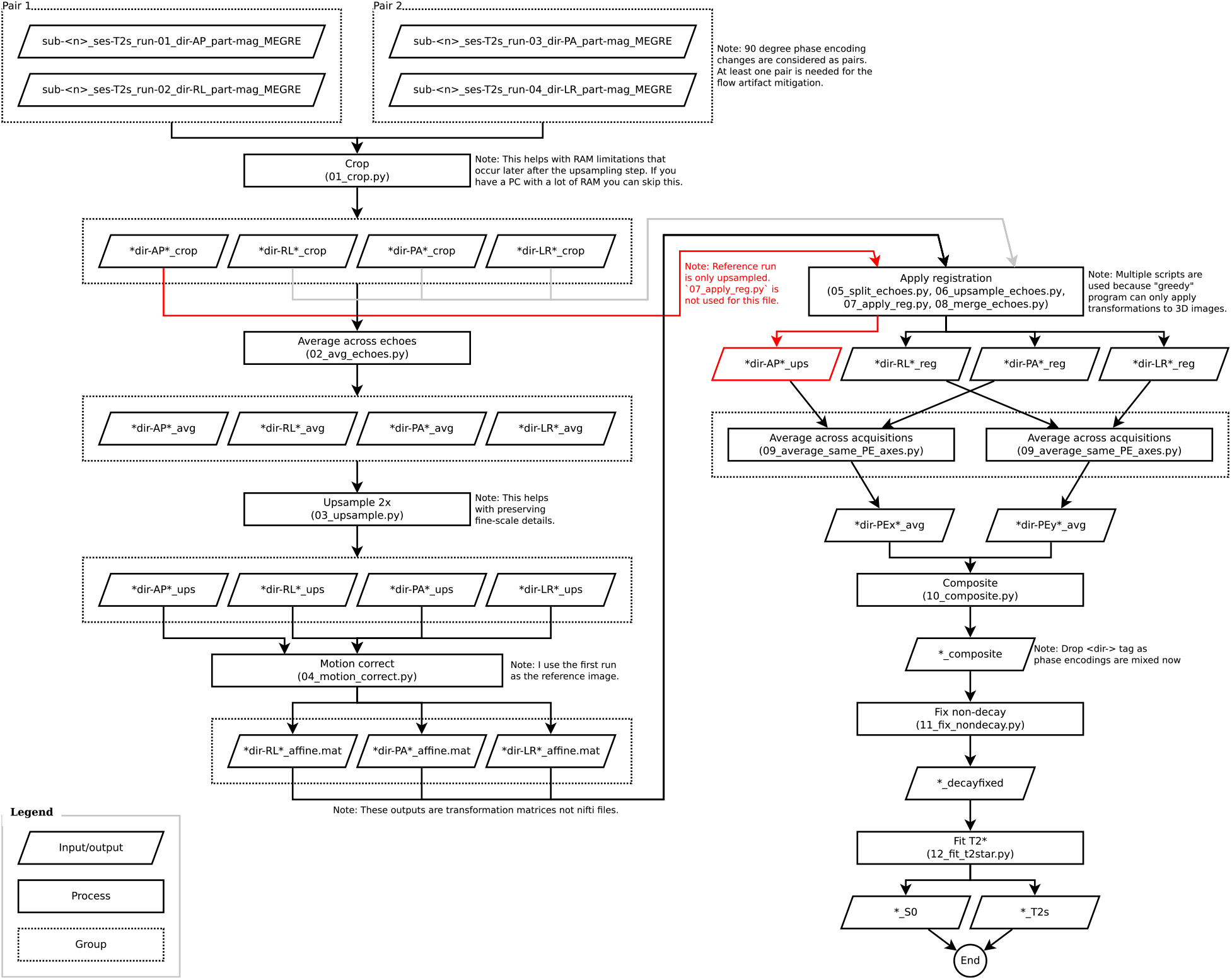
ME GRE flowchart offering a bird’s eye view of our processing steps. Each step refers to a script available at: https://doi.org/10.5281/zenodo.7210342. Note that at the very beginning, we have cropped (01_crop.py) the frontal regions of our images due to computer RAM limitations (32 GB in our case). The frontal regions are cropped because these areas are outside of our regions of interest (calcarine sulcus and Heschl’s Gyrus). This cropping step can be skipped if there is enough RAM available.

**Supplementary Figure 15:**
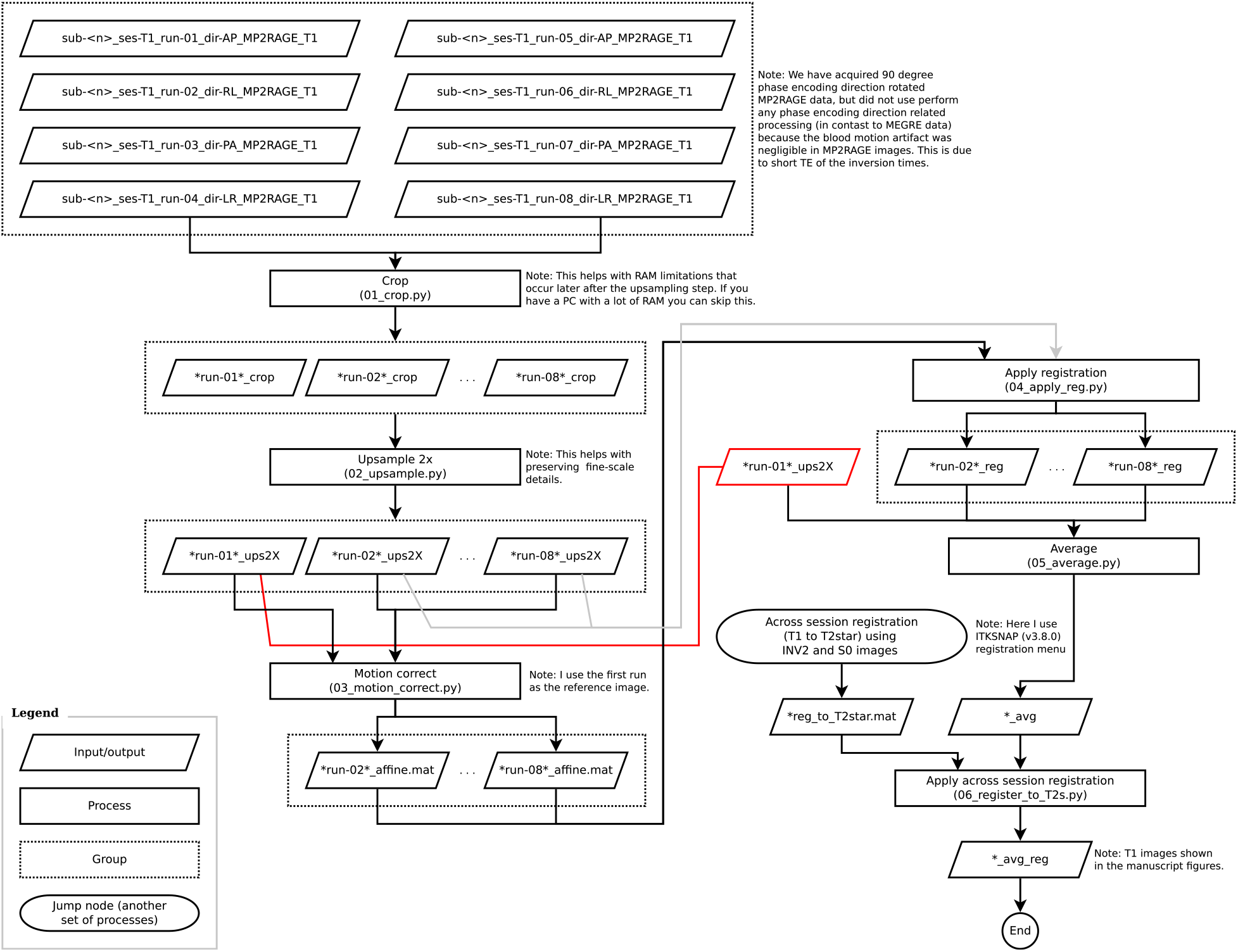
MP2RAGE flowchart offering a bird’s eye view of our processing steps. Format similar to **Supplementary Figure 14**.

**Supplementary Table 1:** Acquisition details of mesoscopic T_2_ and T_2_* contrast MRI datasets. We have strictly selected articles that measure: (i) T_2_ or T_2_* contrasts, (ii) humans, (iii) (in vivo), and (iv) at voxel resolution less than 0.5 mm in at least two dimensions to qualify as “mesoscopic resolution”. Here we summarize acquisition details of these articles (Budde et al., 2011; Duyn et al., 2007; Federau & Gallichan, 2016; Kemper et al., 2018; Lüsebrink et al., 2021; Mattern et al., 2018; Petridou et al., 2010; Sánchez-Panchuelo et al., 2012; Trampel et al., 2011; Zwanenburg et al., 2011). We included Koopmans et al. (2011) as a notable “near-mesoscopic” exception to our criteria (denoted by “*”). Interestingly, most articles did not measure quantitative T_2_* but rather preferred weighted imaging. “3D FatNavs” stands for “fat based motion navigators”, “Preselection” stands for preselecting participants who are experienced in being scanned and can lie still inside the scanner consistently. “N/A” indicates “not reported” or “not applicable”. “?” indicates “cannot be computed given the reported details”.

| Article | Contrast | Pulse Seq. | Voxel size |  |  | Coverage | Quantitative MRI | Motion Correction | Open access to data |
| --- | --- | --- | --- | --- | --- | --- | --- | --- | --- |
|  |  |  | X [mm] | Y [mm] | Z [mm] |  |  |  |  |
| Duyn2007 | $T_2^*$-w | GRE | 0.24 | 0.24 | 1 | Partial | No | N/A | No |
| Petridou2010 | $T_2^*$-w | 2D GRE | 0.25 | 0.25 | 1.5 | Partial | No | Retrospective | No |
| Budde2011 | $T_2^*$ | 3D ME GRE | 0.35 | 0.35 | 2 | Partial | Yes | Retrospective | No |
| Koopmans2011* | $T_2^*$ | 3D ME GRE | 0.75 | 0.75 | 0.75 | Partial | Yes | Retrospective | No |
| Trampel2011 | $T_2$-w | TSE | 0.5 | 0.5 | 0.5 | Partial | No | N/A | No |
| Zwanenburg2011 | $T_2^*$-w | 3D GRE | 0.5 | 0.5 | 0.5 | Partial | No | Retrospective | No |
| Sánchez-Panchuelo2012 | $T_2^*$ | 3D ME GRE | 0.4 | 0.4 | 0.4 | Partial | Yes | N/A | No |
| Federau2016 | $T_2$-w | TSE | 0.38 | 0.38 | 0.38 | Whole brain | No | 3D FatNavs + Retrospective | Yes (https://osf.io/63qjk/) |
| Federau2016 | $T_2^*$-w | GRE | 0.35 | 0.35 | 0.35 | Whole brain | No | 3D FatNavs + Retrospective | Yes (https://osf.io/63qjk/) |
| Mattern2018 | $T_2^*$-w | 3D GRE | 0.33 | 0.33 | 1.25 | Whole brain | No | Prospective + Retrospective | No (but see Lüsebrink2021 for 0.33 mm iso. data) |
| Kemper2018 | $T_2^*$-w | 2D GRE | 0.35 | 0.35 | 0.35 | Partial | No | Retrospective | No |
| Lüsebrink2021 | $T_2$-w | TSE | 0.45 | 0.45 | 0.45 | Whole brain | No | Prospective + Retrospective | Yes (https://doi.org/10.18112/openneuro.ds003563.v1.0.1) |
| Lüsebrink2021 | $T_2^*$-w | GRE | 0.33 | 0.33 | 0.33 | Whole brain | No | Prospective + Retrospective | Yes (https://doi.org/10.18112/openneuro.ds003563.v1.0.1) |
| Gülban2022 | $T_2^*$ | 3D ME GRE | 0.35 | 0.35 | 0.35 | Partial | Yes | Preselection + Retrospective | Yes (https://doi.org/10.17605/OSF.IO/N5BJ7) |

| Article | Contrast | Pulse Seq. | Voxel size |  |  | Voxel volume | Echoes | Slab volume | Parti-cipants | Field strength | Scan duration |  | Total duration | Acceleration |
| --- | --- | --- | --- | --- | --- | --- | --- | --- | --- | --- | --- | --- | --- | --- |
|  |  |  | X [mm] | Y [mm] | Z [mm] |  |  |  |  |  | [min] | [nr. avg.] |  |  |
| Duyn2007 | $T_2^*$-w | GRE | 0.24 | 0.24 | 1 | 0.058 | 1 | ? | 7 | 7 | 10 | 1 | 10 | N/A |
| Petridou2010 | $T_2^*$-w | 2D GRE | 0.25 | 0.25 | 1.5 | 0.094 | 1 | 0.5 | 6 | 7 | 3.6 | 3 | 11 | N/A |
| Budde2011 | $T_2^*$ | 3D ME GRE | 0.35 | 0.35 | 2 | 0.245 | 5 | 0.9 | 5 | 9.4 | 10 | 2 | 20 | GRAPPA = 2 |
| Koopmans2011* | $T_2^*$ | 3D ME GRE | 0.75 | 0.75 | 0.75 | 0.422 | 10 | 0.6 | 8 | 7 | 1.5 | 10 | 15 | GRAPPA = 4 |
| Trampel2011 | $T_2$-w | TSE | 0.5 | 0.5 | 0.5 | 0.125 | 1 | ? | 13 | 7 | 10 | 1 | 10 | Turbo factor = 2 |
| Zwanenburg2011 | $T_2^*$-w | 3D GRE | 0.5 | 0.5 | 0.5 | 0.125 | 1 | 1.5 | 8 | 7 | 6 | 1 | 6 | SENSE = 2.3, Partial Fourier = 0.78 |
| Sánchez-Panchuelo2012 | $T_2^*$ | 3D ME GRE | 0.4 | 0.4 | 0.4 | 0.064 | 4 | 0.9 | 4 | 7 | 15 | 1 | 15 | N/A |
| Federau2016 | $T_2$-w | TSE | 0.38 | 0.38 | 0.38 | 0.055 | 1 | 4.0 | 1 | 7 | 26 | 2 | 52 | PF = 3/4 in both PE dir., Turbo = 166 |
| Federau2016 | $T_2^*$-w | GRE | 0.35 | 0.35 | 0.35 | 0.043 | 1 | 3.1 | 1 | 7 | 42 | 1 | 42 | PF = 3/4 in both PE dir. |
| Mattern2018 | $T_2^*$-w | 3D GRE | 0.33 | 0.33 | 1.25 | 0.136 | 1 | 3.7 | 4 | 7 | 18 | 1 | 18 | N/A |
| Kemper2018 | $T_2^*$-w | 2D GRE | 0.35 | 0.35 | 0.35 | 0.043 | 1 | 0.2 | 2 | 9.4 | 7 | 6 | 42 | N/A |
| Lüsebrink2021 | $T_2$-w | TSE | 0.45 | 0.45 | 0.45 | 0.091 | 1 | 6.3 | 1 | 7 | 23 | 1 | 23 | GRAPPA = 3, PF = 6/8 in Slice dir., Turbo = 173 |
| Lüsebrink2021 | $T_2^*$-w | GRE | 0.33 | 0.33 | 0.33 | 0.036 | 1 | 4.9 | 1 | 7 | 43 | 4 | 172 | PF = 6/8 in PE dir. and in Slice dir. |
| Gülban2022 | $T_2^*$ | 3D ME GRE | 0.35 | 0.35 | 0.35 | 0.043 | 6 | 1.5 | 5 | 7 | 15 | 4 | 60 | GRAPPA = 2 |

